# Topical Formulation of Repurposed FDA-Approved Compounds Inhibits *Pseudomonas aeruginosa* ExoU and Improves Corneal Infection Outcomes

**DOI:** 10.1101/2025.10.26.684634

**Authors:** Daniel M. Foulkes, Keri McLean, Marie Held, John Harris, Yan Sun, Joanne Fothergill, Valerie Price, Dominic P. Byrne, Connie Tam, David G. Fernig, Stephen B. Kaye

## Abstract

Microbial keratitis, infection of the cornea, cause by ExotoxinU (ExoU) strains of *Pseudomonas aeruginosa* have a poor clinical outcome and response to antimicrobials. ExoU, a phospholipase, is secreted directly into host cells, causing their lysis. A screen of 3,034 FDA-approved compounds identified zinc pyrithione (Zp), bismuth subcitrate (Bis), and polymyxin B (Pol) as lead inhibitors of ExoU selectively inhibiting it without affecting human PLA2s or bacterial viability. The compounds have distinct inhibitory mechanisms: Zp disrupted ExoU oligomerization and protein stability; Bis impaired phosphatidylinositol 4,5-bisphosphate-dependent membrane association; Pol directly inhibited catalysis via its lipid-peptide architecture. Compound combinations enhanced ExoU inhibition *in vitro* to nanomolar concentrations. In mammalian cells, Bis and Pol promoted lysosomal trafficking and degradation of ExoU. Using a high-throughput microscopy platform, we screened 53 *P. aeruginosa* keratitis isolates, confirming broad efficacy of these inhibitors in exoU⁺ strains. Therapeutic efficacy was evaluated in *ex vivo* porcine corneas, *Galleria mellonella*, and *in vivo* mouse keratitis models. Topical delivery of ExoU inhibitors, particularly in combination, significantly reduced corneal opacity, ulceration, and stromal damage in porcine corneas without affecting bacterial load. In *Galleria*, compound combinations significantly maintained larval health, improved larval survival and delayed mortality. In a mouse eye infection model, combinatorial treatment reduced disease severity and preserved tissue viability without altering bacterial burden. These findings validate ExoU as a druggable virulence factor and support the repurposing of these compounds as an anti-virulence strategy for the treatment of *P. aeruginosa* infections in humans and veterinary medicine.

## Introduction

The opportunistic pathogen *Pseudomonas aeruginosa* is the primary causative agent of bacterial keratitis and is a leading cause of clinical blindness, particularly in contact lens wearers and immunocompromised individuals [1]. It is also a leading cause of intensive care unit-acquired pneumonia (ICUAP) [2], and is the second most frequent colonising bacteria in patients with COVID19 [2, 3]. As a pathogen of current major concern, the World Health Organisation (WHO) has listed carbapenem-resistant *P. aeruginosa* (CRPA) with the highest priority for the development of new treatments [4].

The type III secretion system (T3SS) is a major virulence determinant in *P. aeruginosa* infections and is strongly associated with poor clinical outcomes, including in pneumonia and microbial keratitis [5, 6]. Among the effectors secreted via this system, ExoU and ExoS are the principal toxins, and their presence is almost always mutually exclusive: strains encoding *exoU* typically lack *exoS* and *vice versa* [7]. ExoU, a patatin-like phospholipase A2, is considered the most cytotoxic T3SS effector and is strongly linked to rapid epithelial injury, tissue necrosis, and severe outcomes such as corneal ulceration and vision loss [8, 9]. In contrast, ExoS is a bifunctional toxin with GTPase-activating and ADP-ribosyltransferase activities, which disrupts host cell signalling and deregulates actin dynamics in more chronic infections [8].

In microbial keratitis, *exoU*^+^ strains predominate, being detected in 61.5% of clinical isolates, and are consistently associated with worse clinical outcomes and increased antimicrobial resistance [10, 11]. Mechanistically, ExoU requires binding to ubiquitin and phosphatidylinositol 4,5-bisphosphate (PIP_2_) for activation [12, 13]. PIP_2_ binding promotes ExoU oligomerisation, markedly enhancing its ubiquitin-dependent catalytic activity. In mammalian cells, ExoU localises to the plasma membrane via its C-terminal four-helical bundle domain, where its phospholipase activity rapidly lyses host cells [14, 15]. This enzymatic activity also liberates arachidonic acid, triggering NF-κB and mitogen-activated protein kinase (MAPK) signalling cascades[16–18]. The subsequent upregulation of proinflammatory mediators, including IL-8 and keratinocyte chemoattractant (KC), drives neutrophil recruitment and intense local inflammation, exacerbating tissue destruction [16, 17]. Collectively, *exoU*^+^ isolates are associated with worse patient outcomes, heightened inflammation, and increased resistance to standard therapies [11].

[10, 19], no ExoU anti-virulence agents have yet been approved for clinical use. Targeting bacterial virulence offers promising therapeutic strategies, with several therapies currently in clinical development [20]. For example, the monoclonal antibody MEDI3902, which targets PcrV of the *P. aeruginosa* T3SS, recently entered Phase II trials for the prevention of ventilator-associated pneumonia but failed to demonstrate efficacy [21]. Traditional antimicrobial agents fail to address ExoU-mediated host cell destruction, and efforts to inhibit the T3SS machinery may risk collateral effects on bacterial physiology and compensatory virulence. Targeting ExoU directly in the host cell, rather than the secretion system, offers a precision approach to anti-virulence therapy, one that could preserve host tissue integrity without promoting resistance or dysbiosis [8, 22].

Repurposing FDA-approved drugs offers a pragmatic strategy for ExoU inhibition, bypassing the lengthy and costly development timelines associated with developing novel small molecules. Here, we employed a high-throughput phospholipase activity screen to identify inhibitors of ExoU from a library of over 3,000 clinically approved compounds. Our screen identified three compounds, zinc pyrithione (Zp), bismuth subcitrate (Bis), and polymyxin B (Pol), that inhibit ExoU via distinct mechanisms, including direct catalytic blockade, conformational destabilisation, and toxin mislocalisation within host cells. Through a series of *in vitro*, cellular, *ex vivo*, and *in vivo* assays, we demonstrate that these agents, especially in combination, significantly reduce ExoU-driven cytotoxicity and preserve epithelial integrity in corneal infection models. These findings establish the feasibility of targeting ExoU directly and lay the foundation for future therapeutic development aimed at mitigating *P. aeruginosa* virulence in ocular and other mucosal infections.

## Materials and methods

The full FDA-approved compound library was purchased from Selleckchem, Houston, USA (3,034 compounds, Catalog No.L1300). Follow up hits for downstream analysis were also purchased separately from Selleckchem. Custom synthesised “Poltide” fatty acid peptide analogues were purchased from Biomatik, Kitchener, Ontario, Canada.

### Recombinant protein production

Full-length *exoU* was cloned into pUCP20T to generate a construct encoding ExoU with a C-terminal 6×His tag. Human phospholipases *PLA2G7* and *PLA2G4C* were cloned into pET28a, each encoding an N-terminal 6×His tag. All plasmid constructs were verified by Sanger sequencing (Eurofins Genomics, Ebersberg, Germany). For protein expression, transformed C43(DE3) *E. coli* were cultured in 8L Terrific broth (Melford Laboratories Ltd, Ipswich, UK) supplemented with ampicillin (for pUCP20T) (100 μg/mL) or kanamycin (for pET41a) (50 μg/mL) and grown to an optical density (OD_600_) of 0.8 before induction of recombinant protein expression using 0.4 mM isopropyl-β-d-thiogalactopyranoside (IPTG). ExoU was expressed for 3 hours at 30°C [22] and Lipoprotein-associated phospholipase A₂ (PLA2G7) and cytosolic PLA₂γ, (PLA2G4C) were expressed overnight at 18 °C. *E. coli* were isolated by centrifugation (5,000 g, 10 minutes) and lysed by sonication in 20 mM Tris-HCl pH 8.2, 300 mM NaCl, 0.1% (v/v) Triton-X-100, 10 mM imidazole, 10% (v/v) glycerol and a complete protease inhibitor cocktail tablet (Roche, Welwyn Garden City, UK). Proteins in the cleared lysate were initially purified immobilised nickel affinity chromatography, followed by size-exclusion chromatography (SEC) (16/600 SuperdexD200, GE Healthcare, Amersham, UK) in 20 mM Tris-HCl pH 8.2, 100 mM NaCl and 10% (v/v) glycerol. Recombinant proteins were frozen in liquid nitrogen and stored at −80°C.

### Phospholipase assays

#### High throughput screening

Screens were performed in 384-well, flat-bottom Corning plates (Corning, Sunderland, UK) in a final volume of 20 µL per well. The reaction buffer was 20 mM Tris-HCl (pHD8.2) and 100 mM NaCl. Each well contained: 175 nM recombinant His₆-ExoU, 5 µM bovine ubiquitin (Sigma-Aldrich, Gillingham, UK), 1 µM PIP₂ (Avanti Polar Lipids, Alabaster, AL, USA), 0.6 mM arachidonoyl thio-phosphatidylcholine (Cambridge Bioscience Ltd., Cambridge, UK), 1 mM DTNB (5,5′-dithiobis-(2-nitrobenzoic acid), Sigma) with DMSO at 0.5% (v/v).

Using an Echo 555 acoustic dispenser, 100DnL of each 10DmM compound stock from the full FDA-approved library (Selleck Chemicals, Houston, TX, USA) was added to assay plates to achieve a final concentration of 50 µM. Reactions were initiated by adding ExoU enzyme, incubated at 25D°C, and absorbance at 414Dnm was measured after 1 h using a Hidex plate reader. Percent inhibition was calculated relative to DMSO controls.

#### IC**₅₀** Determination

Lead compounds were assayed in a 9-point serial dilution series in triplicate. Real-time absorbance readings at 414Dnm were recorded every minute for 60Dmin. For each well, the initial linear portion of the A₄₁₄ vs. time curve (5-15Dmin) was fitted by least-squares regression to yield a slope (ΔA₄₁₄/min). Product formation rates (µmol/min) were calculated using:

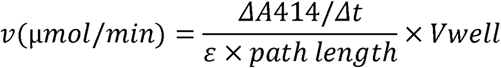

Rates were then normalized to ExoU mass in each well (µg) to give specific activity (µmol/min/µg). Normalized rates were plotted against the logarithm of compound concentration, and IC₅₀ values were determined by fitting a three-parameter nonlinear regression dose-response model in GraphPad Prism.

#### Human phospholipase assays

Recombinant human PLA₂G7 and PLA₂G4C activities were measured in 96-well, UV-transparent microplates (Corning) in a final volume of 50µL per well. Reactions were carried out in 20DmM Tris-HCl (pHD8.2), 100DmM NaCl, and 0.01% TweenD20, with a 1% (v/v) final DMSO concentration. For PLA₂G7, substrate cleavage was monitored using 0.6DmM 2-thio-PAF (Cambridge Bioscience Ltd), and for PLA₂G4C, 0.6DmM arachidonoyl thio-bis-glycerophosphocholine (thio-BGC; Cambridge Bioscience Ltd) was employed. In both assays, 1DmM DTNB (5,5′-dithiobis-(2-nitrobenzoic acid), Sigma-Aldrich) served as the chromogenic reporter. Enzyme (500 nM final) was pre-equilibrated with buffer at 25 °C for 5 min before initiating the reaction by substrate addition. Absorbance at 414 nm was recorded every minute for 1 h on a Hidex plate reader.

#### Secondary Incucyte S3 screen to assess compound efficacy in a wound infection assay

Human corneal epithelial (HCE-T) cells (donated by Kaoru Araki-Sasaki, Japan) were seeded into ImageLock 96-well plates (Sartorius, Epsom, UK) and cultured until confluence in Dulbecco’s modified Eagle’s medium (DMEM)/F-12 medium (Gibco, Thermo Fisher Scientific, Loughborough, UK) with 10% (v/v) foetal bovine serum (FBS; Gibco). Uniform scratch wounds were generated using the IncuCyte WoundMaker (Sartorius) and plates were washed twice with warm phosphate buffered saline (PBS) to remove detached cells.

Cells were infected with *P. aeruginosa* strain PA103 (multiplicity of infection (MOI) 10). In some conditions, moxifloxacin (2.5 µM; 50% minimal inhibitory concentration (MIC)) was added to restrict bacterial overgrowth and extend the assay window [22] [23]. Test compounds were added simultaneously at the following final concentrations: Bis 5 µM, Zp 1 µM, Pol 1 µM, cyclofenil (Cyclo) 5 µM, mivacurium chloride (MivC) 5 µM, and chlorhexidine (Chlorh) 5 µM. All compounds were diluted in DMSO, and vehicle controls (0.1% DMSO) were included in each plate.

To monitor cell viability and lysis, calcein-AM (1 µM; Invitrogen™, Thermo Fisher Scientific, Loughborough, UK) and ethidium homodimer-1 (2 µM; Invitrogen) were added immediately following infection to stain live and membrane-compromised cells respectively. Plates were imaged every 30 minutes for 15 hours using the IncuCyte S3 live-cell imaging system (Sartorius) equipped with a 10x objective. Image acquisition and analysis were performed using IncuCyte S3 software. Quantitative metrics included total live cell area (calcein+) and dead cell count (ethidium+).

Each experimental condition was assayed in duplicate. Post-assay bacterial CFU enumeration was performed in parallel, by serial dilution and agar plating, to confirm that observed cytoprotective effects were not due to antimicrobial activity of the compounds at the concentrations used.

#### Nano Differential Scanning Fluorimetry (nanoDSF)

Thermal stability of recombinant His₆-ExoU was assessed using a Prometheus NT.48 instrument (NanoTemper Technologies, Cambridge, UK) without extrinsic dyes, as these were not compatible with the use of PIP_2_. ExoU was diluted to 5 µM in 20 mM Tris-HCl (pHD8.2), 100 mM NaCl, and 10% (v/v) glycerol. To evaluate ligand effects, four samples were prepared: ExoU alone, ExoU with 5 µM PIP₂, ExoU with 10 µM inhibitor, and ExoU with both 5 µM PIP₂ and 10 µM indicated inhibitor, each maintaining 1% (v/v) DMSO. Ten microliters of each mixture were loaded into high-sensitivity nanoDSF glass capillaries and subjected to a thermal ramp from 25 °C to 95 °C at 1 °C/min. Intrinsic tryptophan fluorescence emission at 330 nm and 350 nm was recorded continuously, and the ratio F₃₅₀/F₃₃₀ was plotted against temperature. Melting temperatures (T⍰) were determined from the inflection point of the first derivative of the fluorescence ratio using the manufacturer’s analysis software. All conditions were measured in triplicate. Changes in melting temperature (Δ T⍰) compared to DMSO, or DMSO PIP_2_ controls were plotted using GraphPad.

#### BS³ crosslinking and SDS-PAGE analysis of ExoU oligomerization

To assess the impact of PIP₂ and small molecule inhibitors on ExoU oligomerization, recombinant His₆ ExoU (5 µM) was first buffer-exchanged into 50DmM HEPES pHD7.4, 100 mM NaCl, 10% (v/v) glycerol using desalting spin columns to remove Tris. Four reaction mixtures (50 µL each) were prepared at room temperature: ExoU alone; ExoU with 5 µM PIP₂; ExoU with 10 µM inhibitor; and ExoU with both 5 µM PIP₂ and 10 µM inhibitor. Bis(sulfosuccinimidyl) suberate (BS³) was added fresh to 1 mM and the samples were incubated for 1 h at 22 °C. Reactions were quenched by addition of Tris HCl pHD7.4 to 50 mM and a further 15 min incubation at room temperature. Each sample was then mixed 1:1 with SDS PAGE loading buffer and resolved on 8% (w/v) Bis Tris SDS-PAGE gels at 200 V. Proteins were transferred to nitrocellulose membranes in Tris-glycine-methanol transfer buffer at 100 V for 1 h at 4 °C. Membranes were blocked in TBST containing 5% (w/v) non-fat milk for 1 h, then incubated overnight at 4 °C with mouse anti-His monoclonal antibody (Bio Rad; 1:2,000). After washing in TBST, membranes were incubated with HRP-conjugated goat anti-mouse IgG for 1 h at room temperature. Signal was detected by chemiluminescence with X ray film exposure, and band patterns were analysed to determine the distribution of monomeric and oligomeric ExoU species under each condition.

#### EGFP-ExoU (S142A) Transfections

HEK293T cells were maintained in Dulbecco’s Modified Eagle Medium (DMEM; Gibco) supplemented with 10% (v/v) FBS, 4 mM L-glutamine, 100 U/mL penicillin and 100 µg/mL streptomycin. The gene encoding the catalytically inactive ExoU S142A was synthesized by GenScript (Piscataway, NJ, USA) and cloned in-frame into the pEGFP-C3 vector using BamHI and NotI restriction sites. Correct insertion and sequence integrity were confirmed by Sanger sequencing (Eurofins Genomics). For analysis of total EGFP-ExoU (S142A) abundance, 2.2D×D10D cells were seeded into 10 cm dishes 24 h prior to transfection. Cells were transfected with 30 µg of pEGFP-C3 vector encoding an N-terminal EGFP-tagged catalytically inactive ExoU (S142A) using polyethylenimine (PEI) (Sigma-Aldrich, Gillingham, UK) at a 3:1 PEI:DNA ratio. After 24 h, cells were lysed and subjected to SDS-PAGE followed by Western blotting using an anti-GFP antibody (Cell Signaling Technology, London, UK) and HRP-conjugated anti-goat secondary antibody. GAPDH was used as a loading control. Protein detection was performed using enhanced chemiluminescence and imaged with a Bio-Rad ChemiDoc imaging system.

#### Confocal Microscopy of EGFP-ExoU and RFP-LAMP1 Co-Localization

HEK293T cells were seeded in 4-compartment glass-bottom imaging dishes (Ibidi, Gräfelfing, Germany) and transfected at approximately 70% confluency with constructs encoding EGFP-ExoU (S142A) (pEGFP-C3 backbone) using polyethylenimine (PEI) at a 3:1 PEI:DNA ratio, and RFP-LAMP1 (CellLight™ Lysosomes-RFP, BacMam 2.0, Thermo Fisher Scientific, Loughborough, UK) by modified baculovirus delivery. Two hours post-transfection, cells were treated with 0.1% (v/v) DMSO or the indicated compounds. Live-cell imaging was performed at 16 h post compound treatment using a Zeiss LSM780 confocal microscope equipped with a 63× oil immersion objective (NA 1.4), a temperature and CO₂ controlled incubation chamber (37°C, 5% (v/v) CO₂), and laser lines at 488 nm and 561 nm for excitation of EGFP and RFP, respectively. Z-stacks were acquired at 0.5 µm intervals with 1024×1024 resolution and a pinhole size of 1 Airy unit.

Colocalisation between EGFP-ExoU(S142A) and lysosomes (Lysosome-RFP, BacMam) was quantified in ImageJ (Fiji) using the Coloc2 plugin. Images were background-subtracted using a 1-pixel rolling ball filter, and Pearson’s correlation coefficients were calculated between the green (ExoU) and red (lysosomal) channels. Data were obtained from three independent experiments.

#### Inhibitor toxicity analysis in human primary corneal cells

Donor human corneas were obtained from the Liverpool Research Eye Biobank (LREB). Corneal rims were quartered and incubated with 1.2 U/mL Dispase II (Roche) in PBS for 2 h at 37 °C. The limbal region was gently scraped using fine forceps to isolate limbal epithelial cells, which were then resuspended and triturated in corneal epithelial cell medium (CECM; DMEM:F12 supplemented with 10% (v/v) FBS, penicillin-streptomycin (100 µg/mL), epidermal growth factor (10 ng/mL), hydrocortisone (0.4Dmg/mL), insulin (5Dmg/mL), adenine (0.18 mM), transferrin (5Dmg/mL), triiodothyronine (T3, 2 nM), and cholera toxin (0.1 nM; Sigma-Aldrich)). Cells were seeded onto mitotically inactivated 3T3-J2 feeder layers, pretreated with 4Dmg/mL mitomycin C for 2 h. Cultures were maintained in CECM, with medium changes three times per week. After 12-14 days of growth, epithelial cells were passaged using 0.5% trypsin-EDTA (Invitrogen) and transferred to 96-well plates for compound cytotoxicity assays.

To assess the cytotoxicity of candidate inhibitors (Bis, Zp, and Pol), lactate dehydrogenase (LDH) release was measured in primary human limbal epithelial cells following 48 hours of compound exposure. Cells were seeded into 96-well plates at a density of 8,000 cells/well in CECM and incubated overnight at 37D°C to allow attachment. After 48 hours of treatment with compounds diluted in CECM, LDH release into the culture supernatant was quantified using the Cytotoxicity Detection Kit (Thermo Fisher) according to the manufacturer’s protocol. Absorbance was measured at 490 nm with a reference wavelength of 620 nm using a Hidex microplate reader. Background signal was subtracted from all wells, and values were normalized to the maximal LDH release control provided in the kit. Each condition was tested in duplicate wells, and the experiment was repeated independently three times (nD=D3). IC₅₀ values were calculated by fitting the data to a nonlinear regression model (inhibitor vs. response, variable slope) using GraphPad Prism.

#### Tertiary CellDiscoverer 7 high-content microscopy screen of clinical *P.***_J***aeruginosa* keratitis isolates

HCE-T cells were seeded at 8,000 cells/well into clear-bottom 96-well plates (Corning). Plates were incubated at 37D°C, 5% (v/v) CO₂ and monitored until cultures reached ∼80% confluence (∼48 h). Immediately prior to infection, DMEM F12 was replaced with DMEM F12 containing 1 µM calcein-AM and 2 µM ethidium homodimer-1 and incubated for 30 min at 37 °C. A panel of 52 clinical *P.*L*aeruginosa* keratitis isolates (28D*exoU*^+^; 24D*exoS*^+^) was used. Bacteria were grown to mid-log phase in LB, washed, and resuspended in PBS. Cells were infected at MOID10 in the presence of Bis 10 µM, Zp 2.5 µM, or Pol 1 µM or indicated combinations thereof. All compounds were prepared in DMSO and added to each well at 0.1% v/v DMSO; vehicle controls received 0.1% DMSO.

Plates were transferred to a Zeiss CellDiscovererD7 (CD7, Carl Zeiss Microscopy GmbH, Germany) equipped with environmental control (37 °C, 5% (v/v) CO₂) and imaged at 5× magnification (0.25 NA). Three-channel images (transmitted light (oblique), Calcein AM and ethidium homodimer-1) were captured automatically at 30 minute intervals over 5 h (Table 1). A bespoke image analysis pipeline developed by Dr. Marie Held at the University of Liverpool Centre for Cell Imaging (CCI) was used. This automated workflow, implemented in Zen Blue and Python, is freely available via Jupyter Notebook. All raw image data are deposited in the EMBL-EBI BioImage Archive.

**Table 1.**
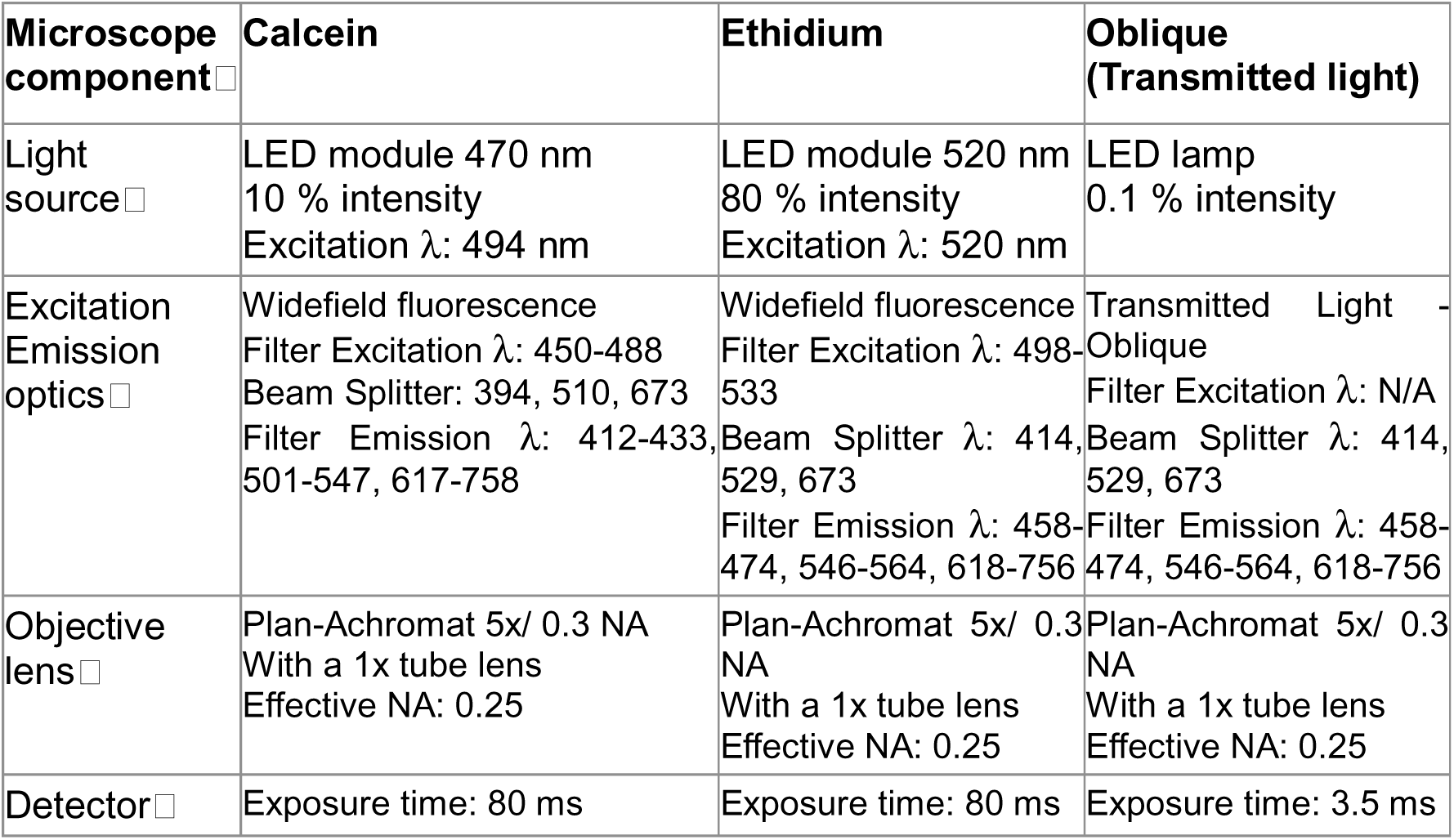
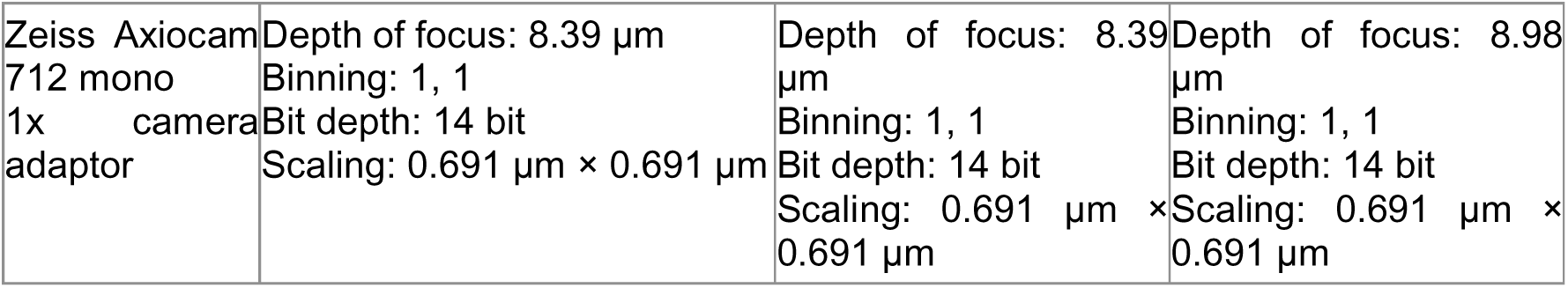
Imaging parameters for live/dead cell microscopy using the Zeiss **CellDiscoverer 7**: Detailed imaging settings used for automated time-lapse acquisition of transmitted light, Calcein AM, and ethidium homodimer-1 signals over 8 hours. The table lists excitation and emission filter specifications, objective and numerical aperture, detectors, and light source parameters for each imaging channel.

| Microscope component | Calcein | Ethidium | Oblique (Transmitted light) |
| --- | --- | --- | --- |
| Light source | LED module 470 nm<br>10 % intensity<br>Excitation λ: 494 nm | LED module 520 nm<br>80 % intensity<br>Excitation λ: 520 nm | LED lamp<br>0.1 % intensity |
| Excitation Emission optics | Widefield fluorescence<br>Filter Excitation λ: 450-488<br>Beam Splitter: 394, 510, 673<br>Filter Emission λ: 412-433, 501-547, 617-758 | Widefield fluorescence<br>Filter Excitation λ: 498-533<br>Beam Splitter λ: 414, 529, 673<br>Filter Emission λ: 458-474, 546-564, 618-756 | Transmitted Light - Oblique<br>Filter Excitation λ: N/A<br>Beam Splitter λ: 414, 529, 673<br>Filter Emission λ: 458-474, 546-564, 618-756 |
| Objective lens | Plan-Achromat 5x/ 0.3 NA<br>With a 1x tube lens<br>Effective NA: 0.25 | Plan-Achromat 5x/ 0.3 NA<br>With a 1x tube lens<br>Effective NA: 0.25 | Plan-Achromat 5x/ 0.3 NA<br>With a 1x tube lens<br>Effective NA: 0.25 |
| Detector | Exposure time: 80 ms | Exposure time: 80 ms | Exposure time: 3.5 ms |
| Zeiss Axiocam<br>712 mono<br>1x camera<br>adaptor | Depth of focus: 8.39 $\mu\text{m}$<br>Binning: 1, 1<br>Bit depth: 14 bit<br>Scaling: 0.691 $\mu\text{m} \times 0.691 \mu\text{m}$ | Depth of focus: 8.39<br>$\mu\text{m}$<br>Binning: 1, 1<br>Bit depth: 14 bit<br>Scaling: 0.691 $\mu\text{m} \times$<br>0.691 $\mu\text{m}$ | Depth of focus: 8.98<br>$\mu\text{m}$<br>Binning: 1, 1<br>Bit depth: 14 bit<br>Scaling: 0.691 $\mu\text{m} \times$<br>0.691 $\mu\text{m}$ |

Images were pre-processed and analysed in Zen Blue (3.7.5, Zeiss, Germany) using an automated macro (<u>Github link to Zen macro</u>). The image was split into the fluorescence channels and the oblique channel. The fluorescence channels were processed via background subtraction (radius = 30) and then merged again with the raw oblique channel. Segmentation was performed in Zen Blue (<u>Github link to</u> <u>analysis settings</u>) using the built-in multichannel segmentation algorithm with background subtraction and thresholding (Otsu method) applied to identify regions of interest (ROIs). Objects were filtered based on size, intensity, and shape metrics to ensure accurate quantification. A minimum object area of 50 µm² and a maximum of 1000 µm² were applied to exclude small artifacts and excessively large structures. Intensity-based filtering was performed on Calcein and EtHD1 fluorescence channels, where objects with mean intensity below 500 (Calcein) or above 2000 (EtHD1) were excluded to remove low-signal noise and overexposed regions. Additional morphological constraints were applied, including convexity, circularity, and compactness thresholds, to refine object selection and remove irregularly shaped debris. Objects overlapping by more than 60% were separated using a watershed-based approach to prevent merging of distinct structures. The final dataset included extracted features such as object count, mean area, relative image area coverage, and integrated density. Image processing and measurements were conducted uniformly across all images the Zen macro which calls the Zen analysis pipeline automatically.

Downstream processing of the measured features, e.g., population average and live/dead ratio calculations were performed in batch in Python, segmented live (calcein⁺) and dead (ethidium⁺) cells and computed total live-cell area (µm²) for assessment of ExoU mediated cell lysis. Cell convexity measurements were performed to assess the impact of ExoS mediated cell rounding. Each condition was assayed in duplicate with data plotted as meanD±DSD of live-cell area (µm²) at 5 h. Figure generation was performed in Python and GraphPad Prism.

#### Quantitative RT-PCR analysis of T3SS gene expression

*P. aeruginosa* strains PA103 and PAO1 were grown overnight in LB at 37 °C with shaking in the presence of Bis 10 µM, Zp 2.5 µM, or Pol 1 µM. EGTA (2.5 mM) was included as a positive control to induce T3SS expression.

Total RNA was extracted from cultures using the RNeasy Mini Kit (Qiagen) according to the manufacturer’s instructions, followed by DNase I treatment to remove residual genomic DNA. First-strand cDNA synthesis was performed using a Reverse Transcription kit (Promega). Quantitative PCR was carried out using SYBR Green Master Mix (Bio-Rad) on an Applied Biosystems StepOnePlus Real-Time PCR instrument.

As previously described, predesigned gene-specific primers were used to amplify *exsA*, *pcrV*, and *exoU* in PA103, and *exsA*, *pcrV*, and *exoS* in PAO1 [22] [23]. RNA polymerase β-subunit gene (rpoB) was used as the internal reference for normalisation. Relative gene expression levels were calculated using the ΔΔCt method, and fold changes were expressed relative to vehicle-treated controls. All reactions were performed in technical triplicates from three independent cultures.

#### *Ex vivo* porcine corneal infection studies

Porcine corneas were processed and infected as previously described [24]. Briefly, porcine eyes were obtained from a local abattoir (CSDMorphet & Sons, Widnes, UK) within 6 h of slaughter and transported on ice. All procedures were conducted under sterile conditions, as described [24]. Corneas with a 2-3 mm scleral rim were excised, immersed in 10 % (v/v) iodinated povidone for 2 min, then rinsed twice in PBS and blotted dry. A single flange of a sterile plumbing ring (B&Q, UK: inner diameter 10 mm) was coated with LIQUIBAND topical skin adhesive and pressed centrally onto the epithelial surface.

To support the stroma and endothelium, 0.5% (w/v) UltraPure agarose in DMEM was prepared by dissolving at 65.5°C, cooling to ∼37°C, and pipetting 1 mL onto the endothelial side. Once solidified, each cornea was transferred, epithelial side up, into a 12IZwell plate. A defined epithelial wound was generated by applying 10 µL 70% ethanol (v/v in PBS) into the ring (corneal apex) for 10Ds, then gently debriding the treated area with a scalpel blade. Corneas were rinsed three times in PBS to remove debris and residual ethanol, and wells were filled with 1 mL DMEM, ensuring the epithelium remained hydrated without submerging the ring.

PA103 was cultured to an OD_600_ of 0.8 in LB, washed twice in PBS, and adjusted to 2D×D10DDCFU/mL. A 50 µL bacterial inoculum (1D×D10DDCFU) was combined with inhibitors Bis 5 µM, Zp 2.5 µM, Pol 1 µM, alone or in combination, or 20 µM moxifloxacin (positive control), all in 0.1% DMSO. Each treatment (or vehicle) was added into the ring, and corneas were incubated at 37°C and 5% CO₂ for 48 h.

Corneal opacity was measured by scanning whole mounts at 400 nm on a Hidex microplate reader and expressed as optical density (OD₄₀₀). Epithelial ulceration was quantified by photographing corneas on a BioIZRad ChemiDoc system and tracing wound areas in ImageJ.

For bacterial burden, rings and adhesive were removed, corneas rinsed in PBS, homogenized in 1 mL PBS using a tissue disruptor, and serial dilutions plated on LB agar. Plates were incubated overnight at 37 °C, and CFU per cornea were determined.

For histology, corneas were fixed in 10 % (v/v) neutralIZbuffered formalin for 24 h, dehydrated, and embedded in paraffin. FiveIZmicron sections were stained with hematoxylin and eosin and scanned on a VENTANA DPD200 slide scanner (Roche).

All conditions were tested in triplicate using corneas from at least three different animals. Data are presented as meanD±DSD. Statistical analyses were performed by oneIZway ANOVA with Tukey’s post hoc test.

### Galleria mellonella infection studies

To assess the therapeutic potential of ExoU inhibitors *in vivo*, *Galleria mellonella* larvae (Pets at Home, UK) weighing 250-300 mg were used. Larvae were stored at 15D°C in the dark and acclimated to room temperature prior to experimentation. Infections were carried out using PA103, grown to mid-log phase in LB, washed twice in PBS, and adjusted to 1D×D10DDCFU/mL. Using a 10 μL Hamilton syringe (Hamilton 702 N, 30 gauge), larvae were injected with 10 μL of bacterial suspension (1D×D10DDCFU) into the last left proleg. Control larvae were injected with 10 μL PBS.

Immediately following infection, larvae were treated topically at the injection site with individual or combined ExoU inhibitors: Bis 0.1 mg/g, Zp 0.01 mg/g, or Pol 0.004 mg/g, alone or in combination. Inhibitors were prepared in PBS containing 0.1% (v/v) DMSO, and 10 μL of each treatment was applied using a micropipette directly to the injection site. DMSO-only vehicle controls were included.

Larvae were incubated in the dark and monitored at 8 h intervals over 40 h. Health was assessed every 8 hours using a validated scoring system [25] which evaluates activity, melanisation, responsiveness to touch, and cocoon formation (maximum score = 10). Since the infection period in our assays was limited to 40 h, no cocoon formation was observed in either infected or control larvae. Consequently, cocoon formation (1.5 points) was excluded from the scoring, and the maximum achievable score in our experiments was 8.5. In parallel, survival was recorded for Kaplan-Meier survival analysis [26].

Clinical score data were analysed using a linear mixed-effects model with repeated measures and Tukey’s post hoc test for multiple comparisons. Survival differences were evaluated using the Kaplan-Meier method with log-rank testing.

### *In vivo* mouse keratitis studies

A previously established scratch infection model was employed in C57BL/6J mice [27]. Female mice (10-12 weeks old; Jackson Laboratory) were anesthetised via intraperitoneal injection (50 μL/25g body weight) of ketamine (50 mg/kg body weight) and dexmedetomidine (0.375 mg/kg body weight), followed by atipamezole (3.75 mg/kg body weight) for reversal. Ethiqa XR (extended-release buprenorphine) was administered to alleviate potential discomfort associated with infection.

A clinical *P. aeruginosa* keratitis isolate strain, PA48386, was cultured in LB broth overnight and then subcultured and grown to log phase (OD_600_ 0.8). PA48386 cells were collected by centrifugation (5,000 g for 5 min) and resuspended in PBS to yield 2 × 10D CFU/mL suspension. On anesthetised C57BL/6J mice corneal infections were induced by creating three parallel epithelial scratches in the central cornea using a 25-gauge needle, followed by topical application of 5 μL PA48386 in PBS (inoculum containing 1D×D10DDCFU). After 30 minutes, mice were treated topically with 5 μL of individual ExoU inhibitors or combinations: Bis 50 μM, Zp 20 μM, Pol 1 μM, or vehicle control (PBS with 0.1% v/v DMSO). Treatments were repeated at 4 h and 24 h post-infection.

Corneal images were collected using a stereomicroscope (Amscope, SM-2TZ) equipped with a 10MP Aptina MT9J003 CMOS colored digital camera at 24 h and 48 h under anaesthesia, after which animals were euthanised for downstream analyses. Corneal disease was scored independently by two masked observers on two separate occasions using a 5-point extended grading system (0-4) previously described [27–29]. This system evaluates four pathological characteristics: (1) area of opacity, (2) density of central opacity, (3) density of peripheral opacity, and (4) epithelial surface quality. Scores for each characteristic were summed, giving a total score ranging from 0 (no infection) to 16 (severe infection). Intra- and inter-observer repeatability were assessed using Cohen’s weighted kappa (SPSS v25). Data were analysed using a general linear model (GLM) with clinical grade on days 1 and 2 as the dependent variable and experimental condition as the factor. A post hoc Tukey test was applied, with significance set at p < 0.05.

### Flow cytometry

Mouse corneas were processed for flow cytometry analysis and CFU enumeration as previously described [29]. Mouse corneas were dissected and digested in collagenase type I (82 U/cornea in 150 μL PBS; Millipore Sigma) at 37 °C for 2 hours. A portion of the digest (5 μL) was serially diluted and plated on LB agar for CFU enumeration. The remaining suspension was processed for flow cytometry to assess immune cell infiltration and viability. Cells were washed and incubated on ice for 10 min in 100 μl ice cold FACS buffer (PBS with 1% FBS) containing 2 μg Fc blocker (anti-mouse CD16/CD32 antibody clone 93; Biolegend). Cell surface staining was conducted on ice for 1 h with APC/Cy7 anti-mouse CD45 (clone 30-F11; Biolegend), PE anti-mouse neutrophil Ly6B.2 (clone 7/4; Abcam), PE/Cy5 anti-mouse F4/80 (clone BM8; Biolegend) antibodies. All samples were also stained with Zombie Violet fixable dye (Biolegend) for dead cells. Stained cells were washed twice with FACS buffer and resuspended in 1% PFA, then detected by BD LSRFortessa flow cytometer. Cells were gated by forward and side scatter, viability, CD45^+^ population, and Ly6B.2^+^ and F4/80^+^ subpopulations. Unstained samples and fluorescence minus one (FMO) control were used to set boundaries for background and positive populations. Analysis of flow cytometric data was performed using BD FlowJo software (v.10). Neutrophils were defined as CD45⁺Ly6B.2⁺F4/80⁻, and non-immune cells as CD45⁻.

### Statistics

Results were obtained from at least three independent experiments unless otherwise stated and are presented as meanD±Dstandard deviation (SD). Statistical analyses were performed using GraphPad Prism and SPSS (version 31). Where appropriate, one-way ANOVA was used to compare multiple groups, followed by Tukey’s post hoc test with correction for multiple comparisons. Significance was defined as *<0.05, **<0.01, ***<0.001, ****<0.0001.

### Ethics

Primary human corneal epithelial cells were derived from donor corneas obtained through the (Liverpool Research Eye Bank with ethical approval (IRAS #239185). Porcine eyes were obtained from CS. Morphet & Sons abattoir (Widnes, UK) as by-products of routine commercial slaughter; no pigs were sacrificed specifically for this study. Work with *Galleria mellonella* does not currently require regulatory approval in the UK. All mouse experiments were conducted under protocols approved by the Cleveland Clinic Institutional Biosafety Committee (IBC protocol 1419) and the Institutional Animal Care and Use Committee (IACUC protocol 0000-2324), in compliance with the Public Health Service (PHS) Policy on Humane Care and Use of Laboratory Animals, the National Institutes of Health (NIH) Office of Laboratory Animal Welfare guidelines, and the Association for Research in Vision and Ophthalmology (ARVO) Statement for the use of animals in ophthalmic and vision research. The Cleveland Clinic’s animal care and use program including Animal Facility is fully accredited by the Association for the Assessment and Accreditation of Laboratory Animal Care (AAALAC) International.

## Results

### Repurposing FDA-approved drugs: a high-throughput screen for ExoU phospholipase inhibitors

Recombinant ExoU bearing an N-terminal 6×His tag was expressed in *E. coli* and purified using immobilised metal affinity chromatography (IMAC) followed by size-exclusion chromatography, as described previously [22] (Supplementary Figure 1A and B). ExoU *in vitro* phospholipase activity was analysed using a real-time, high-throughput phospholipase assay based on a modified version of the Cayman Chemical PLA₂ kit, adapted for 96- and 384-well formats [22] (Supplementary Figure 1C). The assay was conducted in the presence of PIP₂ and ubiquitin, cofactors essential for ExoU activation and catalysis.

In the initial screen of 3,042 FDA-approved compounds from the Selleckchem library 175 nM ExoU was used, which gave a specific activity of 5.85D±D0.26Dμmol/min/μg. Reactions were incubated for 1 h, and absorbance was read at 414 nm (Figure 1A). Compounds that reduced ExoU activity to 40% or less, relative to DMSO-treated controls, were advanced to dose-response analysis. These were performed to eliminate false positives and determine IC₅₀ values of confirmed inhibitors (Figure 1B), with compound structures illustrated in Figure 1C.

**Figure 1.**
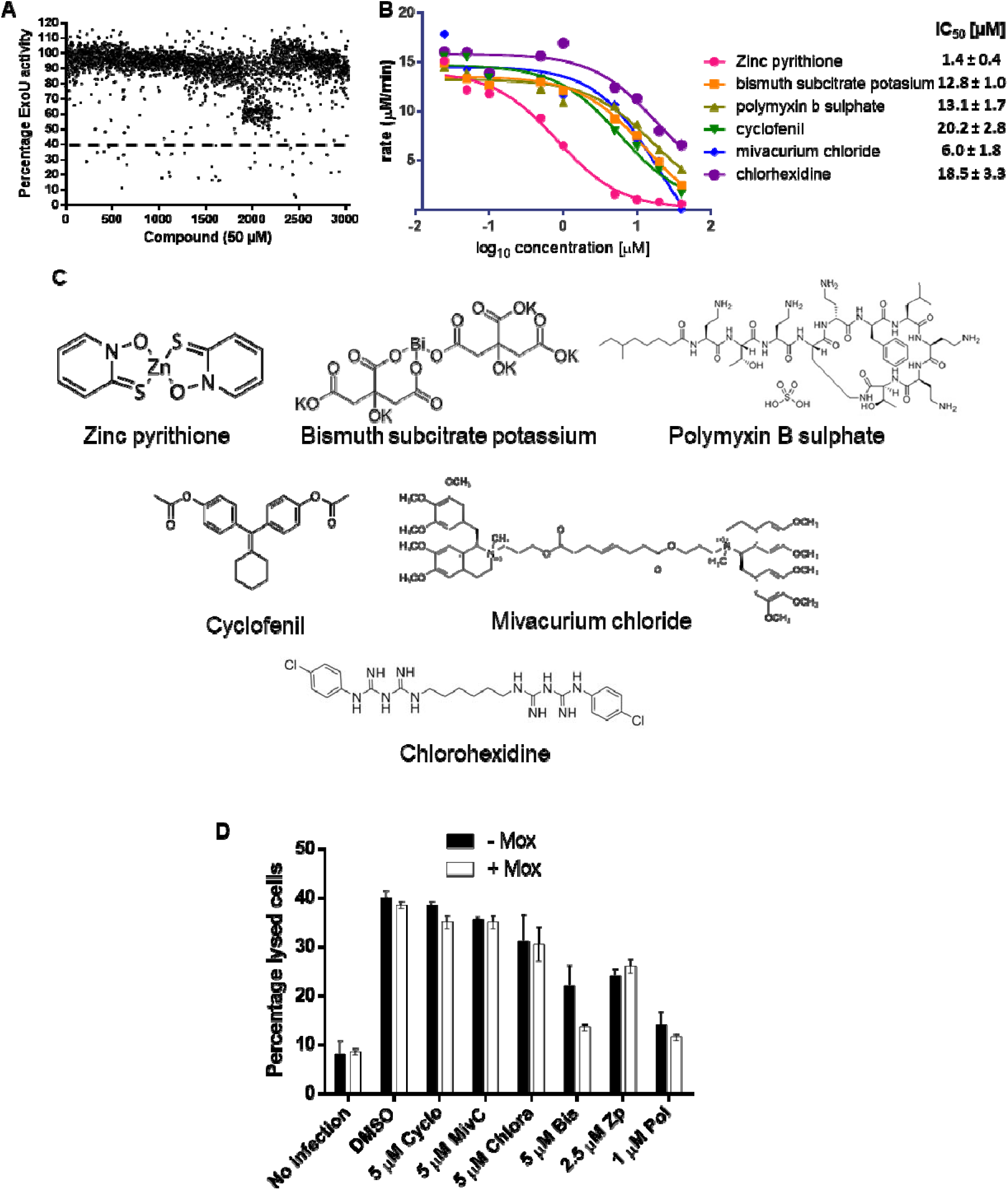
High-throughput screening identifies FDA-approved ExoU inhibitors with activity in vitro and in cell-based infection models. **(A)** A high-throughput *in vitro* phospholipase assay was used to screen 3,042 compounds (50 µM) from the Selleckchem FDA-approved drug library to identify novel inhibitors of ExoU. The graph shows percentage inhibition of ExoU phospholipase activity after a 1 h enzymatic reaction. **(B)** Dose-response curves of selected compounds were generated to determine IC₅₀ values for ExoU inhibition *in vitro*. (**C**) Chemical structures of lead compounds. **(D)** High-content imaging screen of lead ExoU inhibitors using a high-throughput scratch and infection assay. Human corneal epithelial (HCE-t) cells were infected with *P. aeruginosa* PA103 (MOI 10). After 4 hours, cell lysis was quantified using ethidium homodimer staining. The graph shows the percentage of lysed cells as a measure of ExoU cytotoxic activity in the presence of compounds cyclofenil (Cyclo), mivacurium chloride (MivC), chlorhexidine (Chlora), bismuth subcitrate (Bis), zinc pyrithione (Zp), and polymyxin B (Pol).

Of the 35 initial screening hits, we first excluded any compounds that either absorbed at ∼414 nm or otherwise interfered with the DTNB secondary reporter reaction or were unsuitable for therapeutic development (e.g., chemotherapy agents). Six showed clear inhibitory activity in the low micromolar range. Zp emerged as the most potent inhibitor identified to date, with an IC₅₀ of 1.4D±D0.4DμM (Figure 1B) and a Ki of 2.4 μM (Supplementary Figure 2). MivC (IC₅₀: 6.0 ± 1.8 μM, Bis (IC₅₀: 12.8D±D1.0DμM), Pol (IC₅₀: 13.1D±D1.7DμM), Chlorh (IC₅₀: 18.5 ± 3.3 μM), and Cyclo (IC₅₀: 20.2 ± 2.8 μM) were also identified as strong to moderate inhibitors.

With the exception of Cyclo, a selective estrogen receptor modulator with reported adverse effects [30] that make it unsuitable for ocular application, the other hits represent attractive candidates for repurposing as treatments for *P. aeruginosa* keratitis. Notably, several compounds, including Pol and Chlorh are already used topically in ophthalmology [1, 31], enhancing their translational potential. Zp, although not currently used for eye infections is most famously used as an active ingredient in anti-dandruff shampoos. Its applications span across medical treatments (anti-fungal), household products, and industrial materials [32]. Zp exhibits promising potency against ExoU, prompting us to investigate it.

### Determining efficacy of lead ExoU inhibitors in a cellular wound infection model

To evaluate the efficacy of ExoU inhibitors in a cellular context a high-throughput epithelial infection model was developed based on our previously reported scratch wound assay in HCE-T corneal epithelial cells [22]. The assay was adapted to a 96-well plate format compatible with high-content live-cell imaging using the IncuCyte S3 system. Uniform scratch wounds were generated using the IncuCyte WoundMaker, and HCE-T cells were infected with ExoU-producing *P. aeruginosa* PA103 (MOI 10). Cell viability and lysis were continuously monitored for 15 h at 30 min intervals using calcein-AM (viable cell marker) and ethidium homodimer-1 (membrane damage/lysis marker) (Figure 1D).

Because *P. aeruginosa* replicates rapidly *in vitro*, we incorporated sub-MIC moxifloxacin (2.5 μM; 50% MIC) to slow bacterial growth, extend the infection window, and enhance detection of ExoU-dependent cytotoxicity [22]. DMSO-treated, uninfected monolayers exhibited minimal background cell death (8.0 ± 1.8% ethidium-positive cells at 10 h). Sub-MIC Moxifloxacin was confirmed to be non-toxic to HCE-T cells, as ethidium homodimer uptake remained unchanged compared to untreated controls (<1% difference at 10 h). In contrast, infection with PA103 in DMSO-treated wells induced robust ExoU-dependent lysis, with ∼40% of cells staining positive for ethidium homodimer at 10 h (Figure 1D). The inclusion of sub-MIC moxifloxacin did not appreciably reduce cytotoxicity (38.8% lysis), confirming that ExoU-mediated epithelial injury occurs even under conditions of partially restricted bacterial growth (Figure 1D and Supplementary Figure 3B).

We next screened six lead ExoU inhibitors Bis (5 μM), Zp, (2.5 μM), Pol (1 μM), Cyclo (5 μM), MivC (5 μM), and Chlorh (5 μM) in the infection model (Figure 1D). Cyclo, MivC, and Chlorh did not reduce ExoU-mediated lysis (38.5 ± 0.2%, 35.8 ± 0.6%, and 31.2 ± 3.9%, respectively), either alone or in combination with moxifloxacin (35.0 ± 0.6%, 34.8 ± 0.7%, and 30.3 ± 2.6%, respectively). In contrast, Bis, Zp, and Pol showed clear cytoprotection: Bis: 22.1 ± 2.6% lysis (no moxifloxacin); 13.2 ± 0.3% (with moxifloxacin). Zp: 24.1 ± 1.3% (no moxifloxacin); 26.0 ± 1.3% (with moxifloxacin). Pol: 13.9 ± 1.8% (no moxifloxacin); 11.4 ± 0.6% (with moxifloxacin).

Importantly, none of the inhibitors displayed bactericidal activity at the concentrations tested, as demonstrated by bacterial growth dose-response assays in LB and DMEM/F-12 (Supplementary Figure 3, A and C) and endpoint CFU enumeration following IncuCyte imaging (Supplementary Figure 3B). Pol exhibited an IC_50_ of 0.75 μM in LB but 2.1 μM in DMEM/F-12 with 10% FBS, likely due to interactions with serum components; similar trends were observed across a panel of clinical keratitis isolates (Supplementary Figure 3C).

Collectively, these findings identified Pol, Bis, and Zp as promising ExoU targeting inhibitors capable of protecting corneal epithelial cells from *P. aeruginosa* cytotoxicity independent of bacterial killing, establishing this infection model as a robust platform for future screening of anti-virulence therapeutics.

### ExoU inhibitors are non-toxic and do not inhibit human PLA_2_ enzymes

To evaluate host cell compatibility, LDH cytotoxicity assays were performed using primary human corneal epithelial cells (Supplementary Figure 4). Bis showed no measurable toxicity at concentrations up to 160 µM (LD₅₀ >160 µM), while Zp and Pol exhibited LD₅₀ values of 32.6 ± 12.6 µM and 42.5 ± 10.6 µM, respectively. Importantly, these cytotoxic thresholds are an order of magnitude higher than the concentrations applied in our infection assays (Bis, 10 µM; Zp, 2.5 µM; Pol, 1 µM).

In counter screens, we included PLA_2_G7 and PLA_2_G4C which represent two distinct major classes of human calcium-independent PLA_2_ [33, 34], to evaluate the selectivity of our hits for bacterial ExoU over endogenous host phospholipases. Using 2-thio-PAF as the substrate for human PLA2G7 we found that neither Bis, Pol, nor Zp inhibited enzymatic activity, whereas darapladib, a clinical PLA₂G7 inhibitor, effectively suppressed it (Supplementary Figure 5A). In contrast, darapladib did not inhibit ExoU activity when arachidonoyl thio-PC was used as the substrate (Supplementary Figure 5B). Similarly, using arachidonoyl thio-phosphatidylcholine (thio-BGPC) as a substrate for PLA2G4C we observed no inhibition by Bis, Pol or Zp (Supplementary Figure 5C). The use of different thio-ester substrates also underscored this selectivity: while darapladib effectively blocked PLA2G7 with its preferred substrate, it had no effect on ExoU. The absence of inhibition by Bis, Pol and Zp of these enzymes suggests that they selectively targeted ExoU without broadly inhibiting human PLA₂ activity, reducing the likelihood of host toxicity.

All phospholipase assays employed the Ellman reaction, in which free thiols liberated during substrate cleavage react with 5,5′-dithiobis-(2-nitrobenzoic acid) (DTNB), resulting in disulfide bond cleavage and generation of the yellow chromophore 2-nitro-5-thiobenzoate (TNB²⁻). This provides a robust, quantitative readout of enzymatic activity. Under these assay conditions, Bis, Pol and Zp completely abolished ExoU-dependent TNB generation without interfering with DTNB chemistry itself, confirming selective inhibition of ExoU enzymatic activity rather than assay artefact.

### ExoU is inhibited by zinc and bismuth ions

To determine the ExoU inhibitory active pharmacophore in Zp, we assessed copper pyrithione, pyrithione alone and Zn^2+^ in *in vitro* phospholipase assays (Figure 2A). Neither copper pyrithione, nor pyrithione inhibited ExoU (Figure 2A), suggesting that ExoU activity was sensitive to zinc. To further probe this, a panel of metal salts was tested and only zinc sulphate inhibited ExoU phospholipase activity (Figure 2B). We therefore next determined whether Zn^2+^ alone could inhibit ExoU-mediated cytotoxicity (Supplementary Figure 6). LDH release assays in PA103-infected HCE-T cells showed that ZnSO₄ (2.5 µM) did not reduce cell lysis compared to the DMSO control (mean lysis: 89% vs. 85%), whereas Zp (2.5 µM), comprising a pyrithione zinc ionophore, significantly reduced cytotoxicity (mean lysis: 55%). These findings indicate that zinc ions required a carrier to enter host cells and inhibit ExoU activity and that the Zn^2+^chelate Zp is itself active

**Figure 2.**
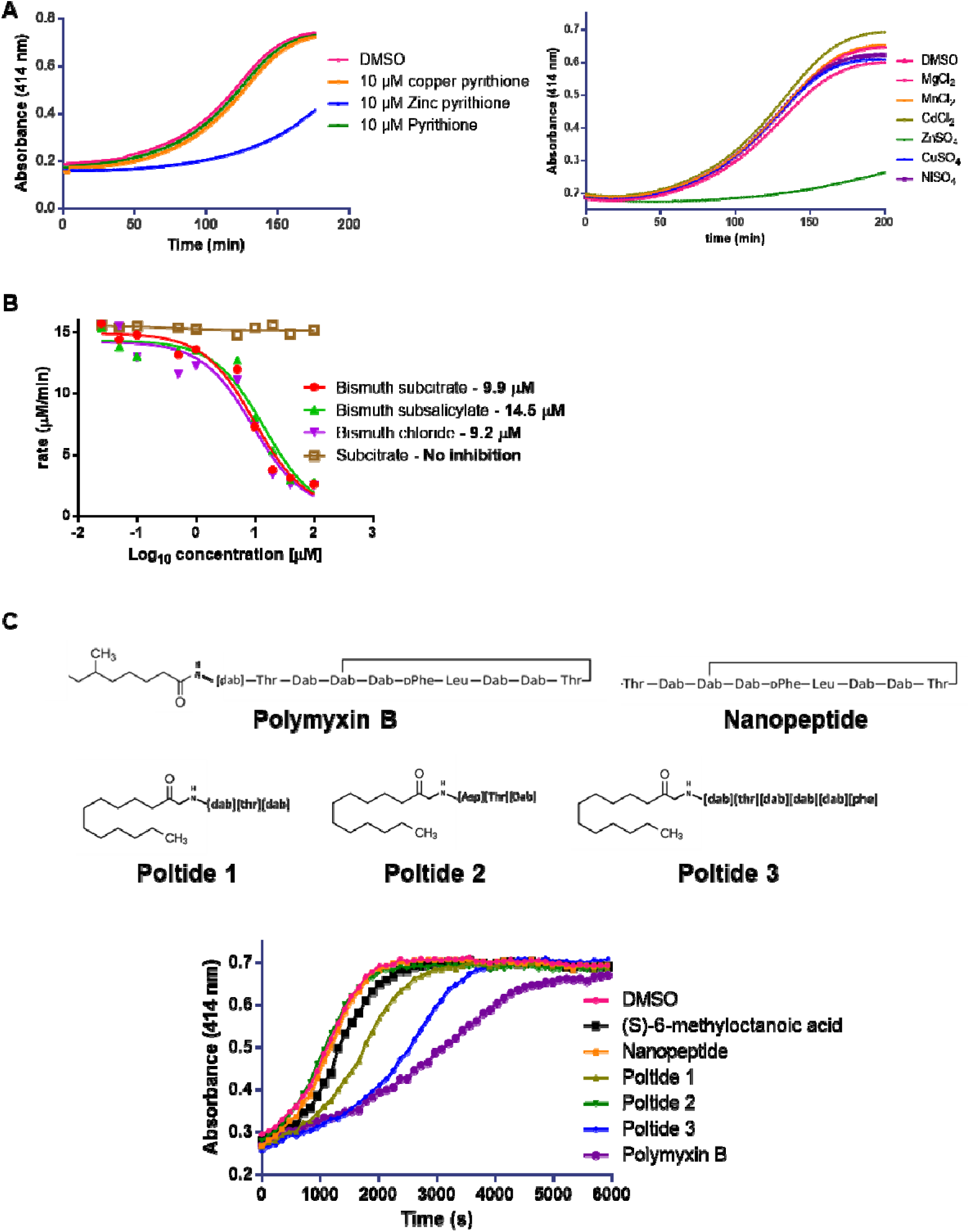
Chemical features associated with ExoU inhibition: metal ions and fatty acid peptides. **(A)** Real-time phospholipase assay showing ExoU inhibition in the presence of pyrithione and pyrithione-coordinated zinc or copper complexes. **(B)** ExoU phospholipase activity detected in the presence of a panel of selected metal salts. **(C)** Dose-response phospholipase assays used to determine IC₅₀ values for bismuth-based ExoU inhibitors and subcitrate alone. **(D)** Chemical structures of native polymyxin B, the cyclic peptide fragment of polymyxin B (nanopeptide), and custom-synthesized polymyxin-derived fatty acid peptides: Poltide 1 (truncated polymyxin mimic), Poltide 2 (negatively charged sequence), and Poltide 3 (linear polymyxin-like analog). **(E)** Real-time phospholipase activity assay demonstrating ExoU inhibitory profiles of each polymyxin compound.

The ExoU inhibitory active pharmacophore in Bis was determined using other bismuth containing compound, bismuth subsalicylate, bismuth chloride and subcitrate in phospholipase assays. Using dose response analysis to elucidate inhibitory IC_50_, Bis (9.9 µM), bismuth subsalicylate (14.5 µM) and bismuth chloride (9.2 µM) inhibited ExoU activity, whereas subcitrate alone did not (Figure 2C).

### Dual-component Inhibition by Polymyxin B

Pol comprises a cyclic cationic decapeptide linked to a hydrophobic fatty acyl tail (Figure 2D) and the inhibitory activity of its structural elements were assayed individually. The isolated hydrophobic tail ((S)-6-methyloctanoic acid) failed to inhibit ExoU (Figure 2E). The cyclic peptide portion of polymyxin B alone (nanopeptide) also did not inhibit ExoU. This indicated to us that Pol’s inhibitory activity likely arose from synergy between its cationic peptide and lipid tail.

The role of the fatty acyl-peptide linkage in ExoU inhibition, was examined by using three custom “poltide” analogues (Figure 2D), each bearing an N-terminal lauric acid (C₁₂) lipid attached to a simplified polymyxin-inspired peptide. Poltide 1 contained an N-terminal lauric acid followed by –Dab–Thr–Dab. In Poltide 2, the charge of the first 2,4-diaminobutyric acid residue was reversed by replacing it with an aspartic acid (poltide 2: lauric acid-Asp-Thr-Dab), while Poltide 3 was extended by consisting of lauric acid followed by Dab–Thr–Dab–Dab–Dab–Phe. Poltide 1 inhibited ExoU, albeit more weakly than Pol (IC₅₀ 40 µM), whereas Poltide 2, bearing an opposite charge in the first amino acid residue, had no effect on ExoU activity (Figure 2E). Poltide 3 was more potent than Poltide 1 (IC₅₀ 20 µM) but did not recapitulate fully the full inhibitory activity of Pol (IC₅₀ 13 µM). These data highlight that both peptide length and charge, as well as the cyclic structure are critical for ExoU inhibition.

### Exploring the *in vitro* biochemical mechanisms of ExoU inhibition by Bis, Pol and Zp

Having prioritized Bis, Pol and Zp as lead ExoU inhibitors, we probed how each agent affected ExoU’s thermal stability and oligomeric assembly using nanoDSF and BS³ crosslinking to dissect weather or not these compounds elicited distinct modes of inhibition.

### Biophysical ExoU inhibitor binding analysis

Nano DSF melt curves (supplementary Figure 7) showed that ExoU alone melted with a midpoint (T⍰) of 40.8 °C. Addition of 5 µM PIP₂ raised the T⍰ to 43.1 °C, confirming that PIP₂ binding stabilizes ExoU’s folded state. As shown in Figure 3A, compound Zp decreased the T⍰ modestly by 1.8 °C without PIP₂ and 1.3 °C with PIP₂, indicating a mild destabilizing effect that was largely independent of lipid. In contrast, Pol produced a substantial destabilization, lowering the T⍰ by 3.8 °C in the absence of PIP₂ and by 6.2 °C in its presence. The disproportionately large ΔT⍰(+PIP₂) for Pol suggests that Pol either competes with PIP₂ for the same binding site or allosterically disrupts the PIP₂-stabilized conformation. Bis had no significant impact on T⍰ in the absence of lipid, but when PIP₂ was present it lowered the T⍰ by ∼2 °C, suggesting that Bis interferes specifically with the PIP₂-ExoU interaction or a PIP₂-induced conformational change.

**Figure 3.**
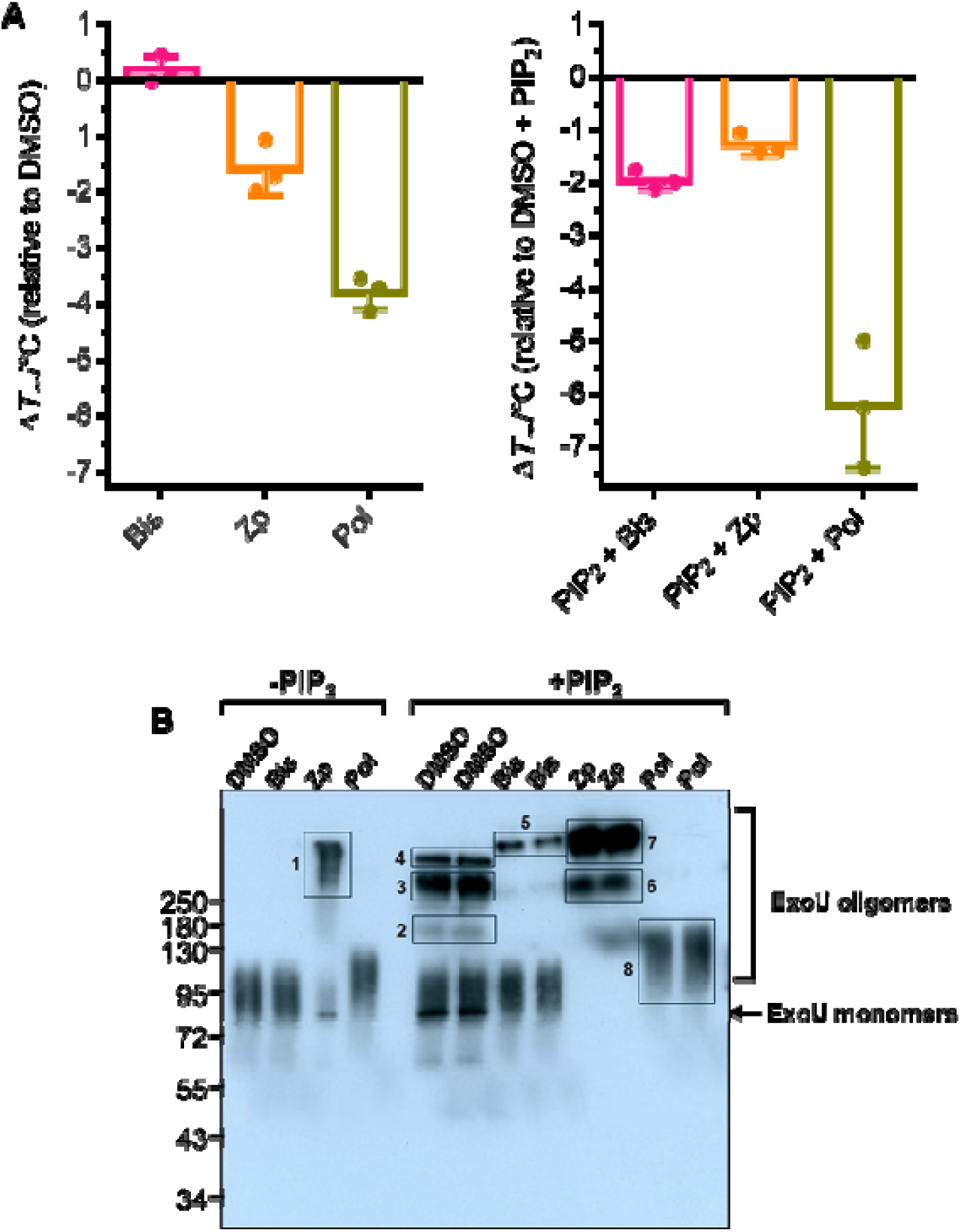
ExoU destabilisation and oligomerisation in the presence of inhibitors and PIP₂. **(A)** Nano differential scanning fluorimetry (nanoDSF) analysis of ExoU thermal stability in the presence of compounds (10 µM) without (left) and with (right) PIP_2_ (5 µM) present. Chart summarises ΔTm values with standard deviation. **(B)** BS³ crosslinking assay followed by SDS-PAGE and anti-6xHis western blot, showing compound-induced changes in ExoU complex formation with and without PIP₂.

Crosslinking of ExoU with BS³, followed by anti-His western blot, provided a more definitive view of oligomeric assemblies under each condition (Figure 3B). In the absence of PIP₂ (lanes 1-4), ExoU alone migrated primarily as a ∼74 kDa monomer with a light smearing, indicating minimal self-association. Addition of Bis mirrored this profile, confirming its inertness in the absence of lipid. However, Zp converted nearly all ExoU into a single, discrete band >250 kDa, with only a faint monomer remaining, while Pol produced a diffuse smear centred around ∼100 kDa, signifying heterogeneous small oligomers.

When PIP₂ was included (Figure 3B, lanes 5-12), ExoU oligomerized robustly in the DMSO control: the dominant species migrated at ∼300 kDa, a ∼400 kDa band was also apparent and a faint ∼150 kDa dimer band was detectable; the ∼74 kDa monomer persisted at moderate intensity with some upward smearing (Figure 3B, lanes 5-6).

In the presence of both Bis and PIP₂ (lanes 7-8), ExoU exhibited broad monomer-region smearing alongside a single discrete high-molecular-weight band matching the Zp (-PIP₂) complex. The absence of lower-order dimers or the 300/400 kDa ladder suggests that Bis constrains PIP₂-induced assembly into one specific oligomeric form. By contrast, Zp + PIP₂ (lanes 9-10) shifted ExoU almost entirely into two uniform bands at ∼300 kDa and >400 kDa, the latter being the most intense species and representing the bulk of crosslinked protein; no monomer remained. Finally, Pol + PIP₂ (lanes 11-12) recapitulated the smear seen without lipid but shifted slightly upward toward the dimer region (∼80-150 kDa), again indicating only modest promotion of heterogeneous, low-order oligomers.

### Bis and Pol destabilise EGFP-tagged ExoU (S142A) expressed in HEK293T cells

To examine the effects of Bis, Pol and Zp on ExoU in host cells, HEK293T cells were transfected with a plasmid encoding EGFP-tagged catalytically inactive ExoU (S142A). As ExoU is toxic to transfected cells, which would preclude analysis due to rapid cell death, the catalytically inactive ExoU (S142A) was used so that measurements could be made 16 h by which time detectable levels of ExoU (S142A) were expressed. Western blotting and confocal fluorescence microscopy were used to establish the effects of the inhibitor (Figure 4).

**Figure 4.**
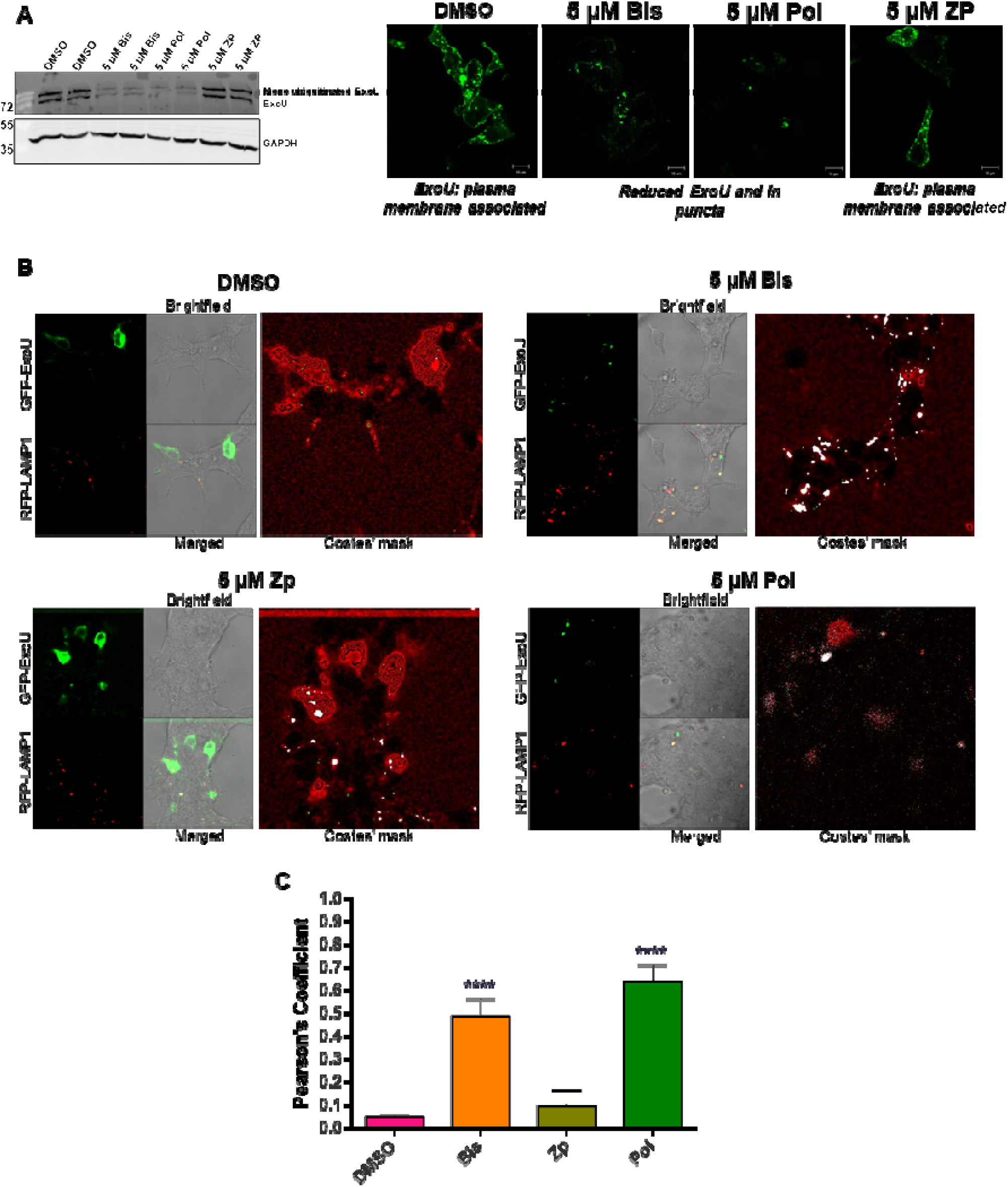
Bismuth subcitrate and polymyxin B promote lysosome localisation and degradation of GFP-S142A ExoU in HEK293T cells. **(A)** Left: HEK293T cells were transfected with pEGFP-C3, encoding S142A ExoU with an N-terminal EGFP tag, in the presence of bismuth subcitrate (Bis), polymyxin B (Pol) or zinc pyrithione (Zp) for 16 hours. Western blotting of whole cell lysates (two independent samples per group; three separate experiments) was used to detect abundance of GFP-S142A ExoU in transfected HEK293T cells, using an α-GFP antibody. Right: Confocal fluorescence microscopy was used to demonstrate plasma membrane distribution of GFP-S142A ExoU in the presence of DMSO and Zp but reduced in the presence of Bis and Pol (full z-stacks are available in supplementary). (**B**) HEK293T cells were co-transfected with GFP-S142A ExoU and the RFP-tagged lysosomal marker LAMP1, followed by 16 h compound treatment. Fluorescence microscopy with split-channel imaging (GFP, RFP, brightfield, and merged), together with Costes’ mask images (colocalised pixels shown in white), was used to visualise S142A ExoU-lysosome colocalization. (**C**) Quantification of ExoU-lysosome colocalization in the presence of DMSO, Bis Zp or Pol was performed from three independent experiments using the Coloc 2 plugin in Fiji (ImageJ) to calculate Pearson’s correlation coefficients. Data were plotted in GraphPad Prism, and statistical significance was assessed by one-way ANOVA with multiple comparisons to DMSO-treated controls.

Western blotting showed that the transfected cells under DMSO treatment expressed consistent quantities of EGFP-tagged ExoU (S142A). The larger form was possibly ubiquitinated, consistent with previous studies, which showed that ExoU was modified by two ubiquitin proteins at Lys178 [35]. The amount of EGFP-ExoU (S142A) in transfected cells treated with either 5 µM Bis or Pol, but not Zp, was sharply reduced (Figure 4A, left panel). Using confocal microscopy, we observed that in the presence of DMSO and Zp, EGFP-ExoU (S142A) was mainly associated with the plasma membrane (Figure 4A, right panel). However, in the presence of Bis or Pol, the level of EGFP-ExoU (S142A) fluorescence was found to be reduced, consistent with the reduction in protein observed in Western blots (Figure 4A, left panel).

We previously demonstrated that ExoU turnover is primarily due to PMSF sensitive proteases rather than the proteosome [23]. Independent studies demonstrated that ExoU(S142A) eventually formed puncta and co-localised with endosomes and lysosomes [35, 36]. We hypothesised that Bis and Pol facilitated ExoU trafficking to and degradation by lysosomes. To assess this, HEK293T cells were we co-transfected with plasmids encoding EGFP-ExoU(S142A) and red fluorescent protein (RFP)-tagged lysosome-associated membrane protein 1 (LAMP1). In DMSO treated cells, 16h post transfection, EGFP-ExoU(S142A) fluorescence remained associated with the plasma membrane (Figure 4B). However, in the presence of Bis or Pol, there was a marked reduction in the amount of EGFP-ExoU(S142A) fluorescence associated with the plasma membrane and the remaining EGFP-ExoU(S142A) formed puncta, co-localising with lysosomes (Figure 4B).

Analysis of the images showed that Bis and Pol treatments (Pearson’s correlation coefficient r=0.49 and r=0.64, respectively) substantially increased colocalisation of ExoU with lysosomes compared with the DMSO control (Pearson’s correlation coefficient r=0.05) (Figure 4C). In contrast, Zp treatment did not enhance colocalisation (Pearson’s correlation coefficient r=0.09). These findings indicated that while all three compounds inhibit ExoU activity *in vitro* and in infection assays, only Bis and Pol promoted trafficking of ExoU to lysosomes within host cells.

### Improved potency of inhibitor combination

Given that Bis, Pol, and Zp inhibited ExoU through distinct mechanisms, we hypothesized that combining these compounds could enhance inhibitory potency. To test this, we performed dose–response phospholipase assays in the presence of Bis, Zp, Pol, and an equimolar combination of all three (Figure 5).

**Figure 5.**
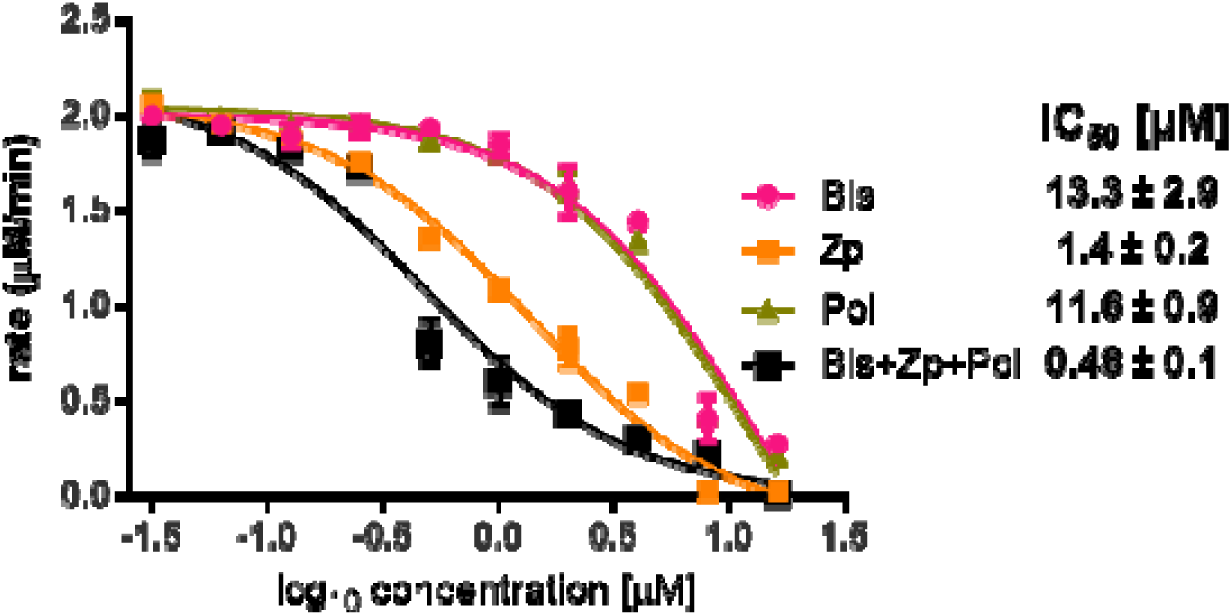
Combinatorial compound treatment enhances ExoU inhibition *in vitro*. Phospholipase activity of ExoU was measured in real-time assays in the presence of individual inhibitors (Bis, Zp, or Pol) or an equimolar combination of all three (Bis+Zp+Pol). Dose-response curves were generated by fitting log[inhibitor] versus activity data using a three-parameter nonlinear regression model in GraphPad Prism. IC_₅₀_ were derived from the fitted curves to quantify the extent of ExoU inhibition.

Individually, Bis, Zp, and Pol inhibited ExoU activity with IC_₅₀_ values of 13.3 ± 2.9 µM, 1.4 ± 0.2 µM, and 11.6 ± 0.9 µM, respectively. When combined, the mixture displayed markedly enhanced potency, yielding an IC_₅₀_ of 0.48 ± 0.1 µM, suggesting that the compounds act through complementary mechanisms, providing rationale for assessment of inhibitor combination treatments in compound efficacy studies.

### Efficacy of ExoU inhibitors in a high content microscopy screen of clinical *P. aeruginosa* keratitis isolates

Following confirmation that Bis, Pol, and Zp reduce ExoU-mediated cytotoxicity in cell infection models using the laboratory PA103 strain, we extended our evaluation to a clinically relevant panel of strains of *P. aeruginosa* using a fully sequenced, phenotyped, and genotyped library of 52 *P. aeruginosa* clinical keratitis isolates from the UK Microbiology Ophthalmic group [37]. High-throughput phenotypic screening used a CellDiscoverer 7 (CD7) platform for live-cell imaging.

HCE-T cells were seeded in 96-well plates, stained with Calcein-AM and ethidium homodimer, and infected at ∼80% confluence with each clinical isolate (MOI 10) in the presence of individual inhibitors (Bis, Pol or Zp) and two examples of the paired combinations (Bis+Zp, Bis+Pol) (Figure 6). Time-lapse imaging was performed every 0.5 h over 4.5 h. Among the 52 isolates, 28 were *exoU*⁺ and 24 *exoS*⁺. For nearly all exoU⁺ strains, ExoU-mediated cytolysis was evident as progressive loss of Calcein-AM fluorescence and membrane rupture. The sole exception was isolate 68228, which displayed fewer lysed cells than uninfected controls (Figure 6B-F), indicating an absence of ExoU-dependent cytotoxicity. In contrast, *exoS*⁺ isolates induced cell rounding while maintaining Calcein-AM signal, consistent with cytoskeletal disruption but preserved membrane integrity.

**Figure 6.**
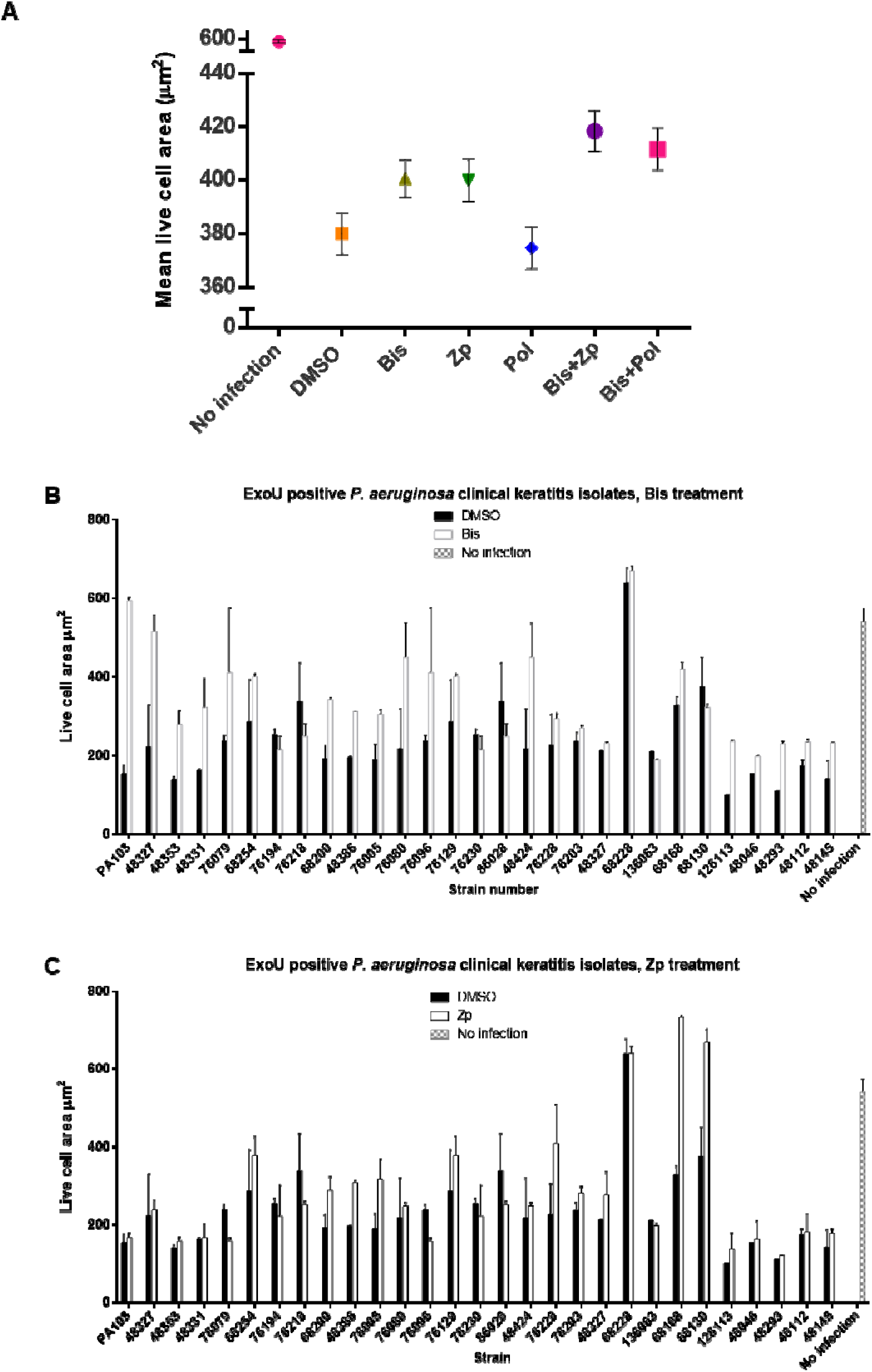
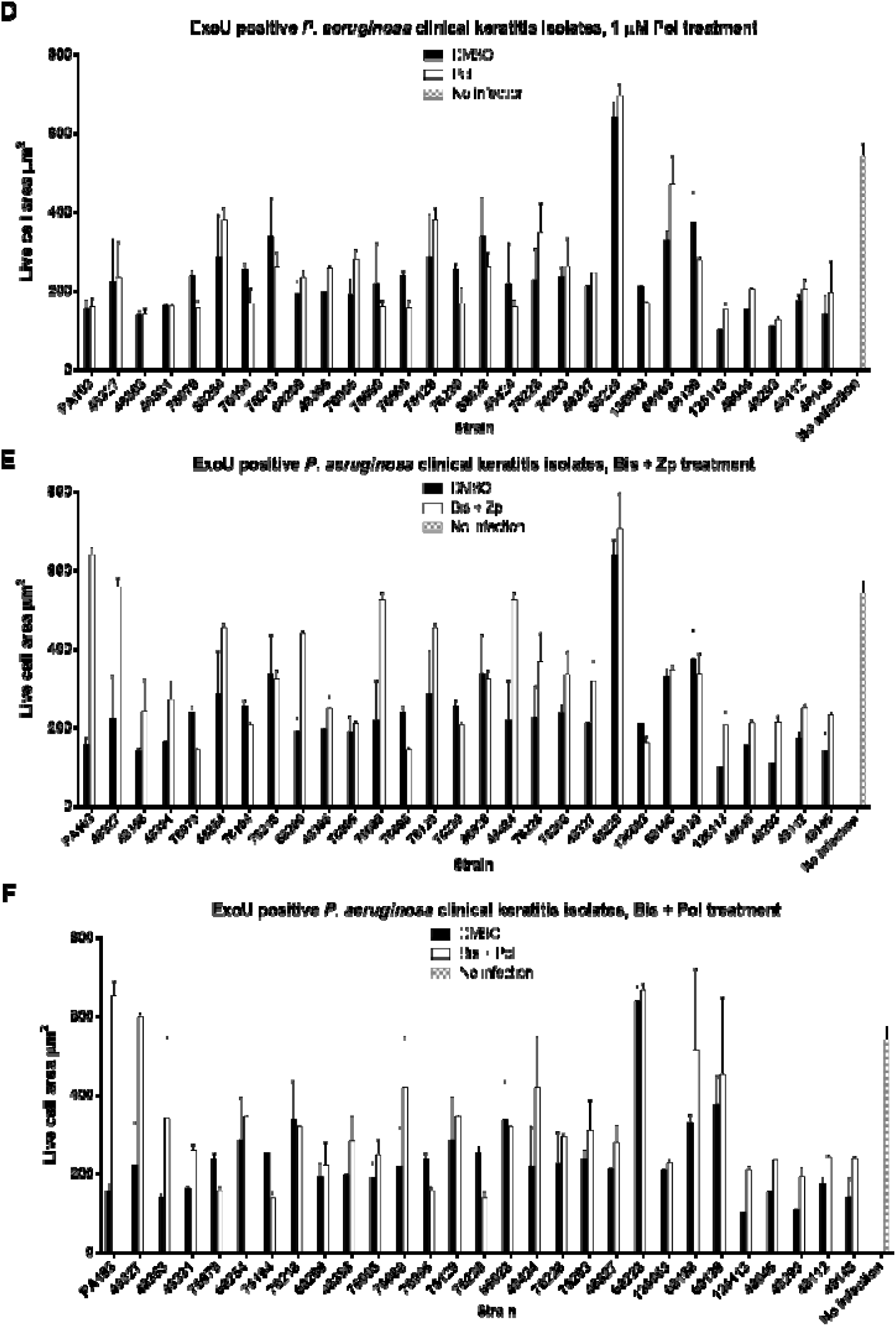
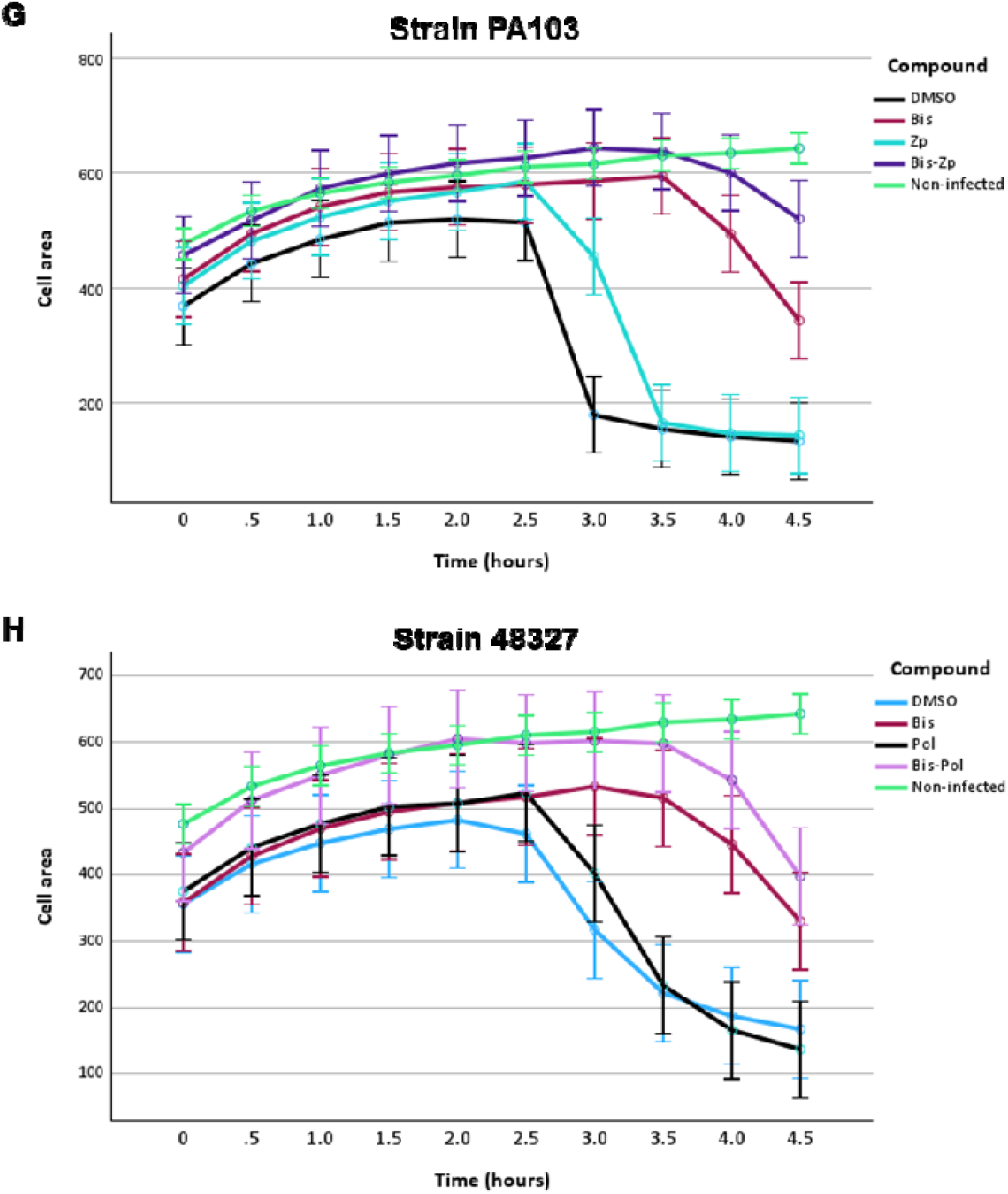
High-content imaging screen to evaluate ExoU inhibitor efficacy across clinical *P. aeruginosa* keratitis isolates. A high-content microscopy screen using the Cell Discoverer 7 (CD7) platform was developed to assess the efficacy of ExoU inhibitors against PA103 and a panel of 29 *exoU^+^ P. aeruginosa* clinical keratitis isolates. HCE-T cells were seeded into 96-well plates and grown to 80% confluence prior to infection at MOI 10. Cells were stained with calcein-AM and ethidium homodimer to assess cell lysis over 4.5 h. Live cell area (µM^2^) was calculated at 30-minute intervals over 4.5 hours of infection. (**A**) Chart showing the calculated mean live cell area for each condition calculated across the entire time series. **(B-F)** Bar charts showing each live cell area at 4.5 h post infection for infection with *ExoU^+^* clinical isolates in the presence of: 10 µM Bis; **(B)** 2.5_JµM zinc pyrithione (Zp); **(C**) 1_µM Pol; **(D)** combination of Bis and Zp; **(E)** combination of Bis and Pol; **(F)**. Uninfected controls are indicated by checkered bars. **(G-H)** Full time series analysis for **(G)** PA103 and **(H)** strain 48327 showing reduced cell lysis with compound combination treatments compared to single compound treatments.

### Compound efficacy in ExoU producing clinical *P. aeruginosa* clinical isolates

In infections with ExoU-producing isolates, all three compounds and combinations conferred measurable protection against cytotoxicity (Figure 6). To capture an overall measure of protection across the entire experiment, the mean live cell area was calculated at each timepoint within each time series and compared treatments (Figure 6A). This analysis revealed significant differences between groups (p=0.02) (Supplementary Table 1), with the Bis+Zp combination providing the strongest overall protection. Specifically, Bis+Zp significantly outperformed DMSO (mean difference 38.36, p < 0.001), Bis (17.73, p < 0.001), Pol (43.66, p < 0.001), and Zp alone (18.42, p < 0.001), and was comparable to Bis-Pol (p = 0.27) (Supplementary Table 1).

At the 4.5 h snapshot, Bis (10 µM) provided measurable protection in 48% of clinical isolates, Zp (2.5 µM) in 52%, and Pol (1 µM) in 48% (Figure 6, B-D). While Bis generally elicited the greatest increase in viable cell area, Zp showed unique activity against isolate 68130, which was unresponsive to other treatments. Notably, the non-responsive subset of strains was otherwise largely conserved across compounds. Combination therapy improved coverage: Bis+Zp protected 66% of isolates, and Bis+Pol protected 62% (Figure 6, E-F). However, six cytotoxic isolates (76194, 76218, 76230, 86028 and 136063) remained refractory to all treatments, showing lysis equivalent to DMSO controls by 4.5 h. These recalcitrant strains may reflect intrinsic differences in ExoU expression levels, host cell interactions, or additional virulence mechanisms. Supplementary Figure 8 provides the full-time course of live-cell area detected across all (*exoU^+^* and *exoS^+^*) clinical keratitis isolates.

Time-series analyses further highlighted strain-specific responses. For PA103, the Bis+Zp combination provided the most durable protection (Figure 6G), whereas for isolate 48327, Bis+Pol was superior (Figure 6H). Taken together, these findings demonstrate a variability of ExoU inhibition across clinical isolates and suggest that, in a clinical setting where strain susceptibility is unknown, combination therapy incorporating Bis, Zp, and Pol would provide the broadest protective benefit.

### Bis, Pol and Zp do not effectively reduce ExoS mediated cytotoxicity

To determine whether our compounds specifically inhibit ExoU rather than broadly interfering with type III secretion, we examined the 24 *exoS*⁺ isolates in our clinical panel and quantified host cell convexity as a readout of ExoS-mediated cytotoxicity (Supplementary Figure 9). ExoS promotes cell rounding through its GAP and ADP-ribosyltransferase domains, which inactivate Rho family GTPases, disrupt actin polymerisation and focal adhesion dynamics, and thereby increase cell convexity [38, 39]. Across the full 4.5 h infection window, none of the compounds or their combinations altered cell convexity compared to DMSO controls (Supplementary Table 2). At the 4.5 h endpoint, Bis treatment modestly reduced convexity in four strains (Supplementary Figure 9A), while Zp impacted only one strain (136083) (Supplementary Figure 9B), Pol had no measurable effect (Supplementary Figure 9C), and the combinations Bis+Zp and Bis+Pol altered convexity in just three and two strains, respectively (Supplementary Figure 9, D-E). Taken together, these observations indicated that while isolated strain-specific differences were detected, Bis, Zp, and Pol generally did not interfere with ExoS-driven cytopathic responses. This supports the assertion that their primary activity is selective inhibition of ExoU.

In addition, we evaluated whether Bis, Zp or Pol affected T3SS gene expression in *P. aeruginosa* PA103 and PAO1 [23]. Quantitative RT-PCR analysis (Supplementary Figure 10) revealed no significant changes in the expression of key T3SS-associated genes *exsA*, *pcrV*, and *exoU* in PA103, or *exsA*, *pcrV*, and *exoS* in PAO1, following treatment with the compounds at concentrations used in our cell-based assays. These results indicate that the compounds do not suppress T3SS gene transcription under these growth conditions, suggesting that any protective effects observed in cell assays are not due to inhibition of T3SS gene expression.

### Evaluation of ExoU inhibitors in a porcine *ex vivo* corneal infection model

Building on our previously developed *ex vivo* porcine corneal infection model [24], which uses medical adhesive and 10 mm plumbing rings to localize both bacterial inoculum and topical treatments, the therapeutic potential of the lead ExoU inhibitors was assessed in this more disaease relevant setting. Using this platform, we evaluated the ability of Bis, Zp and Pol to mitigate *P. aeruginosa* PA103-induced corneal damage. The ring-based setup allowed sustained exposure of dissected porcine corneas to both compound and pathogen at the infection site. We quantified infection severity through measurements of corneal opacity, epithelial ulceration, and bacterial burden (CFU per cornea), and further assessed tissue integrity by histological examination using haematoxylin and eosin (H&E) staining. Dissected porcine corneas were topically infected with 50 µL solutions of P. aeruginosa strain PA103 (1×10D CFU) mixed and treated topically with 50 µL solutions of Bis (5 µM), Zp (2.5 µM), or Pol (1 µM), either alone or in combination. After 48 hours, visual inspection revealed that monotherapies had only modest effects on reducing corneal opacity, with Bis performing slightly better than Zp. Combinatorial treatments yielded progressively improved outcomes: Bis+Zp showed minor improvement over single-agent treatments, while Zp+Pol and Bis+Pol produced greater reductions in opacity. Notably, the triple combination (Bis+Zp+Pol) resulted in corneas that closely resembled uninfected controls, suggesting near-complete protection (Figure 7A).

**Figure 7:**
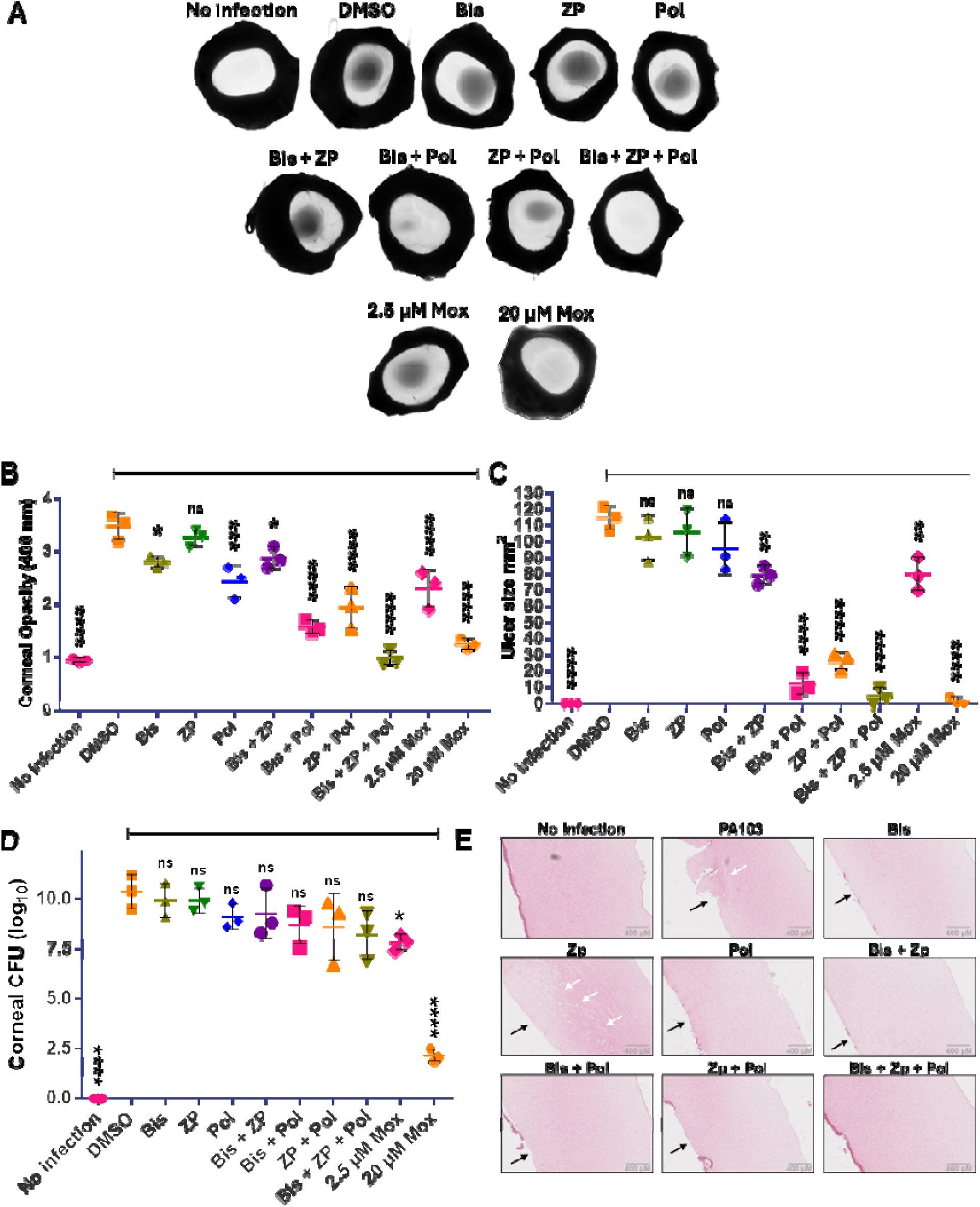
ExoU inhibitor combinations reduce *P. aeruginosa p*athogenicity in *ex vivo* porcine corneas. **(A)** Dissected porcine corneas were infected with 1×10_J CFU of PA103, followed by topical application of ExoU inhibitors individually (Bis: 5 µM, ZP: 2.5 µM, Pol: 1 µM) or in combination. A 20 µM moxifloxacin treatment served as a positive control. Corneas were imaged after 48 hours of infection using a BioRad ChemiDok imaging system. **(B)** Corneal opacity caused by ulceration 48 hours post-infection was quantified by measuring optical density at 400 nm using a spectrophotometer. **(C)** The ulcerated area (mm²) of infected corneas was measured at 48 hours post-infection using ImageJ. **(D)** At 48 hours post-infection, corneas were washed with PBS, homogenized, and serially plated onto agar plates to determine the bacterial load (CFU) within the corneal tissue. **(E)** Infected corneas were fixed in paraformaldehyde (PFA), paraffin-embedded, sectioned, and stained with hematoxylin and eosin (H&E). Sections were imaged using a Ventana DP 200 slide scanner. Black arrows indicate areas of epithelial erosion/ulceration; white arrows highlight regions of stromal edema.

These qualitative observations were supported by quantitative measurement of corneal opacity (OD₄₀₀; Figure 7B). While individual compounds modestly reduced opacity compared to DMSO-treated infected controls, combinations, particularly Bis+Pol and the Bis+Zp+Pol treatment, produced significantly greater improvement. The triple combination was comparable to 20 µM moxifloxacin, a clinically relevant positive control. Assessment of epithelial ulceration (Figure 7C) mirrored these trends. Corneal damage was substantially reduced in combination treatments, with Bis+Pol and Bis+Zp+Pol showing the most dramatic reductions in ulcer size, approaching the resolution achieved with high-dose moxifloxacin. Bacterial loads within the corneas, measured by CFU (Figure 7D), were not significantly altered by compound treatment, indicating that ExoU inhibitors were not directly bactericidal under these conditions. As expected, 20 µM moxifloxacin markedly reduced bacterial burden but did not fully eradicate infection, consistent with previous reports [24].

Histological analysis of H&E-stained sections (Figure 7E) corroborated these findings. DMSO-treated infected corneas showed complete epithelial loss, severe stromal erosion, and edema. In contrast, Bis and Pol treated corneas showed partial epithelial preservation and less stromal damage. The triple combination conferred near-complete epithelial integrity and minimal stromal swelling, closely resembling uninfected tissue.

Together, these data demonstrated that the three ExoU inhibitors, particularly in combination, could significantly reduce the severity of *P. aeruginosa* PA103-induced keratitis in an *ex vivo* porcine model, even without directly affecting bacterial viability.

### Bis, Zp and Pol enhance survival of PA103 infected *Galleria mellonella* larvae

To assess the therapeutic potential of ExoU inhibitors *in vivo*, *Galleria mellonella* larvae were infected with 1×10D CFU of PA103 and treated topically with individual compounds or combinations thereof. Mean clinical health scores, using a validated (0-8.5) scoring system [25] were analysed over 40 h of infection (Figure 8A) by linear mixed models with repeated measures and post hoc Tukey testing (Table 2). Larval health was monitored at 8h intervals over 40 h (Figure 8B) and survival was assessed by Kaplan-Meier analysis (Figure 8C).

**Figure 8:**
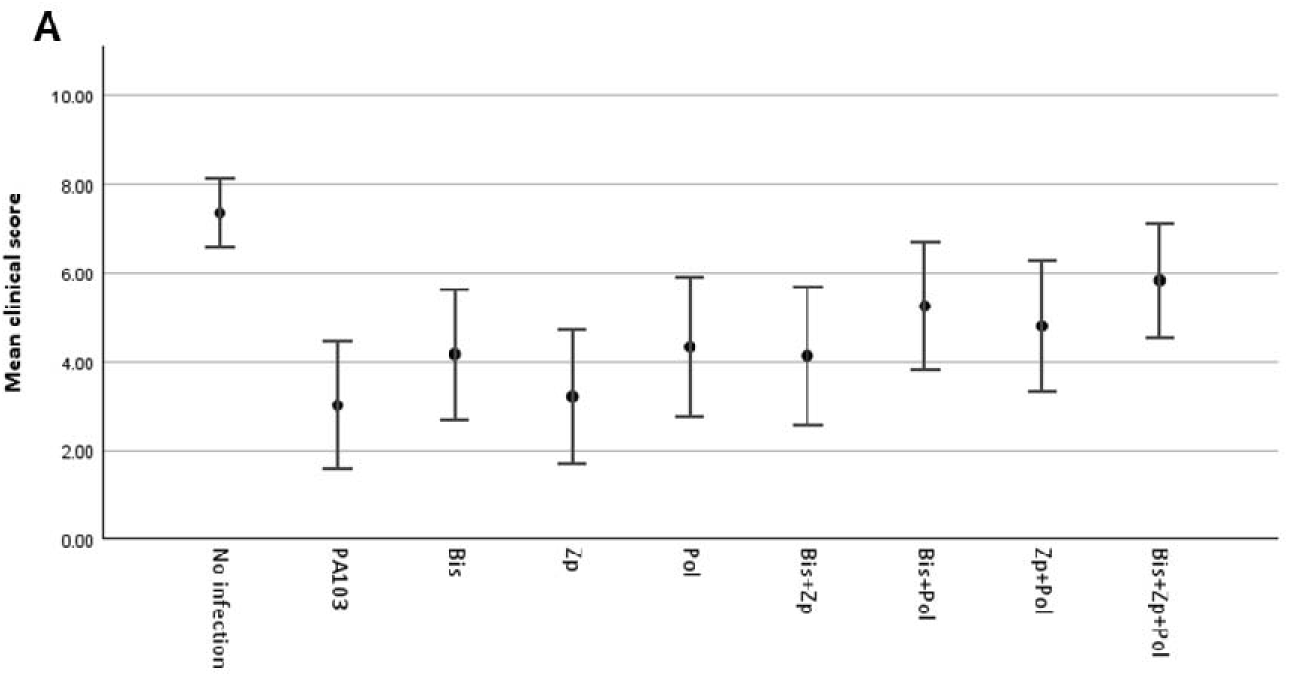
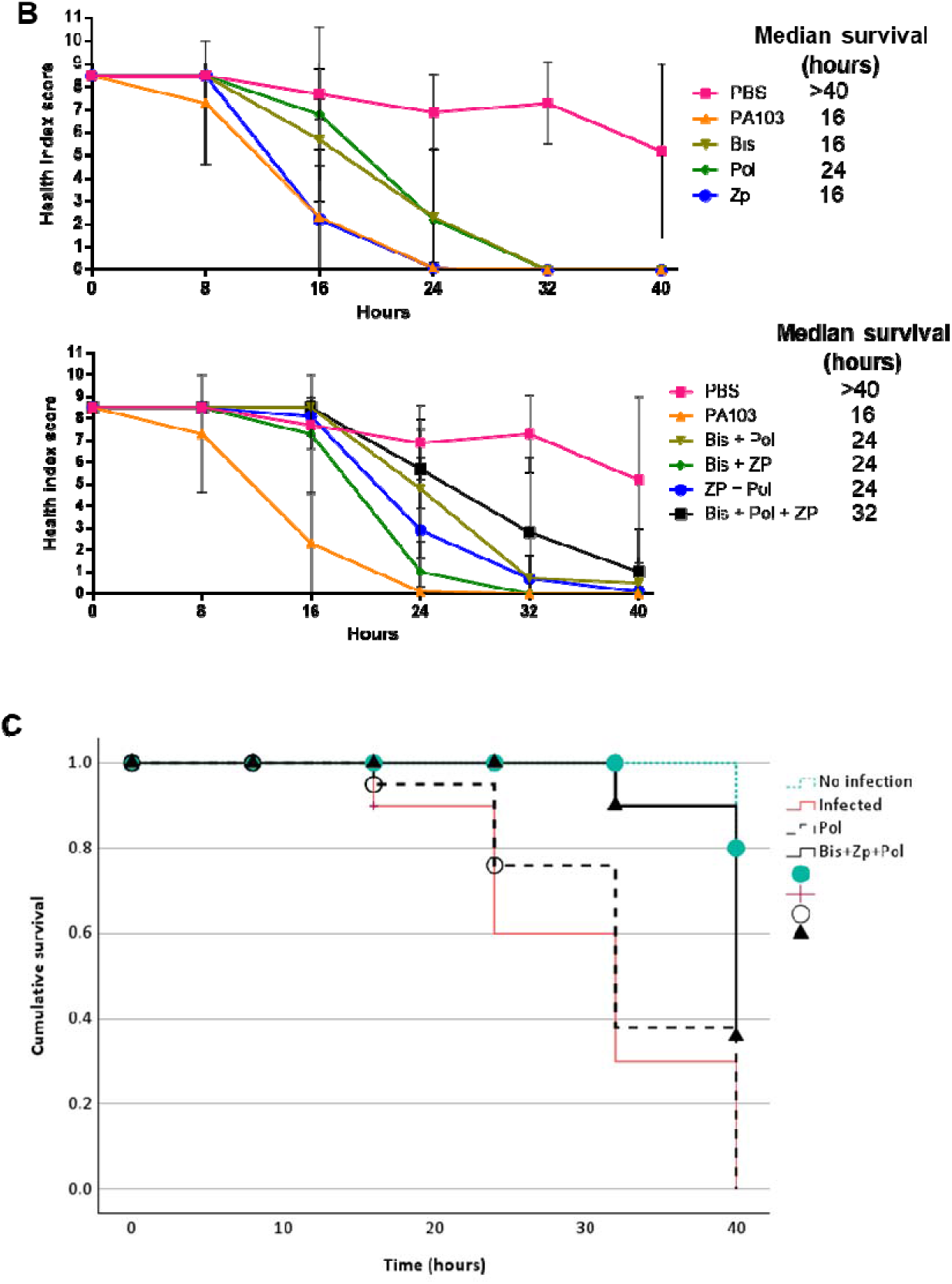
ExoU inhibitors enhance survival of Galleria mellonella infected with PA103. Galleria mellonella larvae were infected with 1 × 10_J CFU of P. aeruginosa strain PA103 and treated with ExoU inhibitors or vehicle control (PBS with 0.1% DMSO). Treatments included bismuth subcitrate (Bis; 0.1 mg/g), zinc pyrithione (Zp; 0.01 mg/g), polymyxin B (Pol; 0.004 mg/g), or combinations thereof. **(A)** Infection progression was monitored over 40 hours using the established *Galleria mellonella* Health Index scoring system. The top panel shows data for individual compound treatments; the bottom panel shows results for compound combinations. **(B)** Kaplan-Meier survival analysis was used to assess the protective effects of ExoU inhibitor treatments. Left: survival curves for single compound treatments. Right: survival outcomes for combination treatments.

**Table 2.** Statistical analysis of mean clinical health scores (5 Galleria larvae per group), using a validated (0-8.5) scoring system, were analysed across 40h of infection, which were analysed by linear mixed models in SPSS with repeated measures and post hoc Tukey testing. Statistical differences between all groups are shown. Those groups between which there was found to be statistically significant differences are marked with an Astrex.

| Galleria | Mean (SD)<br>Clinical score | Condition |  |  |  |  |  |  |  |
| --- | --- | --- | --- | --- | --- | --- | --- | --- | --- |
|  |  | Bis | Zp | Pol | Bis+Zp | Bis+Pol | Zp+Pol | Bis+Zp+Pol | DMSO |
| No infection | 7.35 | *$<0.01$ | *$<0.01$ | *$<0.01$ | *$<0.01$ | *0.02 | *$<0.01$ | 0.27 | *$<0.01$ |
| Bis | 4.17 | | 0.54 | 1.00 | 1.00 | 0.27 | 0.09 | *$<0.01$ | 0.22 |
| Zp | 3.22 | | | 0.49 | 0.70 | *$<0.01$ | 0.04 | *$<0.01$ | 1.00 |
| Pol | 4.33 |  |  |  | 1.0 | 0.70 | 0.99 | 0.10 | 0.23 |
| Bis+Zp | 4.13 |  |  |  |  | 0.38 | 0.91 | *0.02 | 0.39 |
| Bis+Pol | 5.25 | | | | | | 0.99 | 0.96 | *$<0.01$ |
| Zp+Pol | 4.80 | | | | | | | 0.46 | *$<0.01$ |
| Bis+Zp+Pol | 5.83 | | | | | | | | *$<0.01$ |
| Vehicle | 3.21 |  |  |  |  |  |  |  |  |

PBS-injected controls remained healthy throughout the experiment (mean score 7.4 ± 2.0). Infection with PA103 caused rapid health decline, with complete mortality by 32 h (Figure 8, A and B). Compared to uninfected PBS controls, infection significantly reduced clinical scores in mock-treated larvae (PBS + 0.1% DMSO; p<0.01) and in all single-agent (Bis, Zp, Pol) and dual-treatment groups (Bis+Zp, Pol+Zp and Bis+Pol p<0.01). By contrast, the triple combination (Bis+Zp+Pol, mean score 5.8 ± 3.4) did not significantly reduce clinical scores relative to uninfected controls (p=0.27), indicating superior protection (Table 2).

Among single-agent treatments, Bis (0.1 mg/g) and Zp (0.01 mg/g) provided modest, transient protection, delaying mortality by ∼8 h in a subset of larvae. Pol (0.004 mg/g) offered slightly stronger protection, with some larvae surviving beyond 24 h, but none prevented mortality by 40 h (Figure 8B). The dual combinations improved outcomes: Bis+Zp delayed larval decline, though mortality still occurred by 32 h; Zp+Pol extended survival slightly further, with some larvae retaining partial activity up to 40 h. Bis+Pol provided stronger protection, with 1-2 larvae maintaining health scores at 40 h (Figure 8B).

The triple combination (Bis+Zp+Pol) conferred the greatest benefit. All larvae remained healthy through 16 h, several maintained scores ≥4.5 at 24 h, and one larva retained a score of 4.5 at 40 h (Figure 8B). Kaplan-Meier analysis confirmed a significant survival improvement for this combination compared to infection alone, with outcomes not significantly different from uninfected controls (p=0.27; Figure 8C). These findings demonstrated that combining the ExoU inhibitors provided superior *in vivo* protection, with the triple regimen markedly improving survival in this lethal infection model.

### ExoU inhibitor combinations improve clinical outcome in an *in vivo* mouse model of keratitis

To evaluate the therapeutic potential of ExoU inhibitors during *P. aeruginosa* corneal infection, we used a murine scratch and infection model [27, 29]. Mice received topical treatment with individual inhibitors or inhibitor combinations at 30 min, 4 h, and 24 h post-infection. Disease severity was evaluated at 24 and 48 h using a standard 16-point clinical scoring system (Figure 9A), with representative corneal images shown at 24 and 48 h (Figure 9B). Bacterial load was quantified by CFU enumeration (Figure 9C). In parallel, flow cytometric analysis of digested corneas was performed to assess host responses, including corneal cell viability, total leukocyte infiltration, and neutrophil recruitment (Figure 9D-G).

**Figure 9:**
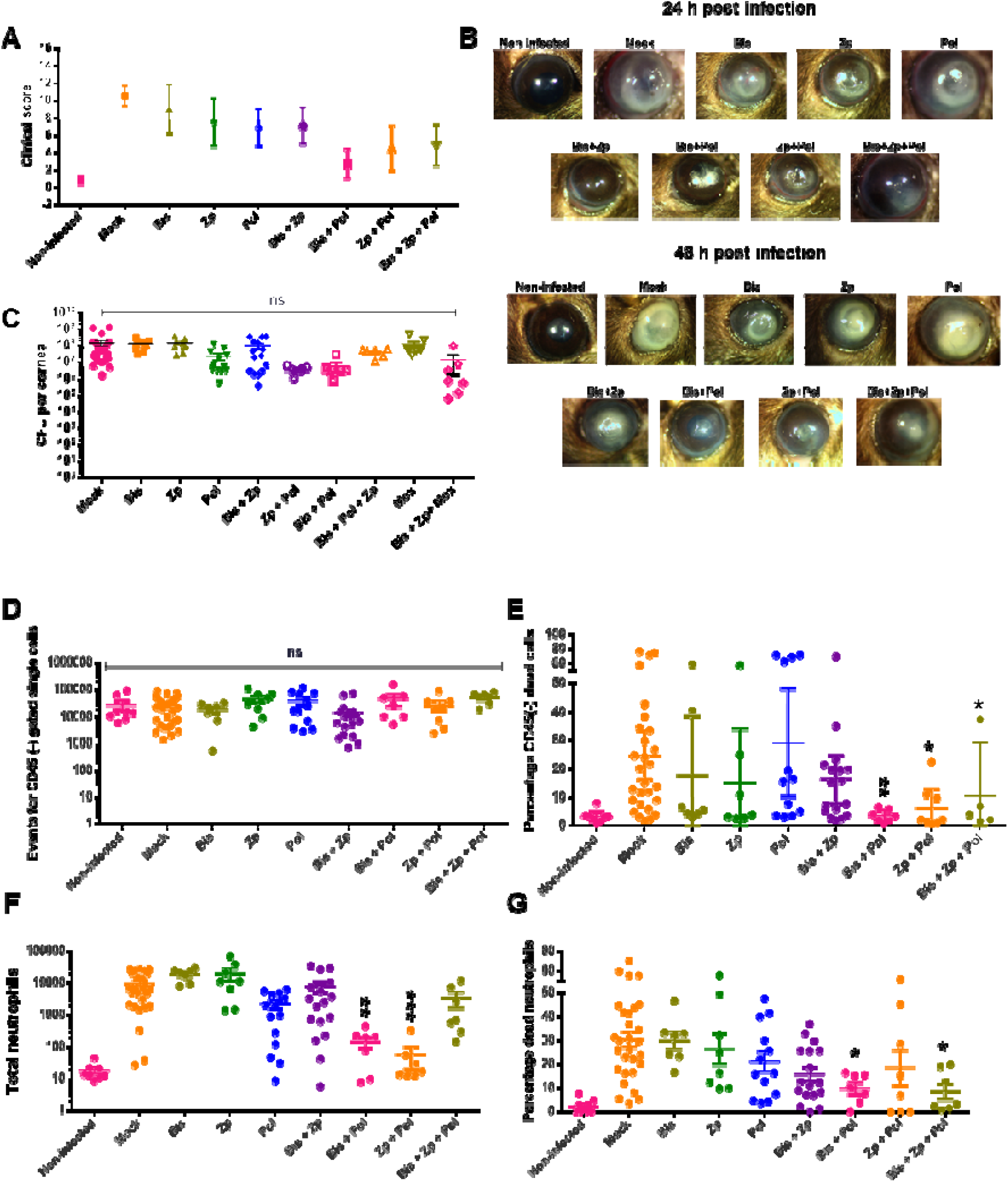
ExoU inhibitor combinations reduce disease severity and bacterial burden in a murine *P. aeruginosa* corneal infection model. Corneal infections were established in female C57BL/6J mice (8–10 weeks old) by creating three parallel scratches with a 25-gauge needle, followed by topical inoculation with 1 × 10_J CFU of *P. aeruginosa* strain PA48386. After 30 min, mice received topical treatment with ExoU inhibitors Bis (50 µM), Zp (20 µM), Pol (1 µM) or PBS vehicle control (5 µL per cornea). Treatments were repeated at 4 h and 24 h post-infection. Eyes were imaged at 24 h and 48 h, and corneas collected at 48 h for analysis. **(A)** Clinical severity of corneal infection at 48 h, scored on a standardized 16-point scale by two independent, blinded assessors. Bar graphs show severity scores. (**B**) representative corneal images are shown below for each treatment at 24 h and 48 h. **(C)** Bacterial burden (CFU) in whole corneal homogenates at 48 h. **(D-G)** Flow cytometry of corneal single-cell suspensions. **(D)** Quantification of non-immune cells (CD45⁻). **(E)** Proportion of dead non-immune cells (Zombie Violet⁺). **(F)** Quantification of neutrophils (CD45⁺Ly6B⁺F4/80⁻). **(G)** Proportion of dead neutrophils across treatment groups. Statistical significance was assessed by one-way ANOVA with post hoc correction for multiple comparisons against mock-treated controls. *p < 0.05, **p < 0.01, ***p < 0.001.

### Assessment of clinical score and analysis of corneal CFU

564 images were taken at 24 and 48 h post inoculation. All images were graded twice each on two separate occasions by 2 specialists blinded to the conditions. Cohen’s weighted kappa for the composite score was 0.84 and 0.77 for intra-rater and 0.74 and 0.78 between raters first and second grading. A General Linear Model (GLM) was used to analyse the results with the clinical grade at 24 h (Table 3) and 48 h (Table 4) as the dependent variables and the different experimental conditions as factors. A post hoc Tukey test was used with p<0.05. There was no significant difference in the clinical score between strains PA103 and PA48386 (p=0.104). The clinical scores for each condition for PA48386 on days 1 and 2 are presented in Tables 3 and 4 and figures 9A and B. The trends for days 1 and 2 were similar but the degree of disease was greater at 24 h (mean 10.73 (SD 4.60)) compared to 48 h (5.65 (2.83)) (p<0.01).

**Table 3:** Clinical severity scores at 24 hours post-infection in a murine *P. aeruginosa* corneal infection model. Female C57BL/6J mice (8–10 weeks old) were scratched and topically inoculated with *P. aeruginosa* strain PA48386 (1 × 10_J CFU per cornea). Mice received topical treatment with ExoU inhibitors Bis (50 µM), Zp (20 µM), Pol (1 µM), or combinations thereof at 30 min, 4 h, and 24 h post-infection. Control animals received PBS with 0.1% (v/v) DMSO. Corneal images were captured 24 h post-infection and graded by two blinded observers using a standardized 16-point scoring system. Values represent mean ± SD from two independent, blinded assessments. Statistical analysis was performed using a General Linear Model (GLM) followed by Tukey’s post hoc test (*p* < 0.05). Combination treatments reduced clinical severity compared to mock treatment, with the lowest scores observed in the Bis + Pol and Zp + Pol groups.

| 24 h | Mean (SD)<br>Clinical<br>score | Condition |  |  |  |  |  |  |  |
| --- | --- | --- | --- | --- | --- | --- | --- | --- | --- |
| Condition |  | Bis | Zp | Pol | Bis+Zp | Bis+Pol | Zp+Pol | Bis+Zp+Pol | Infection |
| No infection | 0.17 (0.35) | p<0.01 | <0.01 | 0.14 | 0.001 | 0.64 | 0.93 | 0.94 | <0.01 |
| Bis | 4.99 (3.60) |  | 0.25 | 0.18 | 0.99 | 0.24 | 0.06 | 0.02 | 0.99 |
| Zp | 7.25(1.96) |  |  | <0.01 | 0.007 | <0.01 | <0.01 | <0.01 | 0.41 |
| Pol | 2.92 (2.46) |  |  |  | 0.43 | 1.00 | 0.95 | 0.64 | <0.01 |
| Bis+Zp | 4.30 (3.08) |  |  |  |  | 0.52 | 0.15 | 0.005 | 0.18 |
| Bis+Pol | 2.45 (2.22) |  |  |  |  |  | 1.0 | 0.99 | 0.007 |
| Zp+Pol | 1.75 (2.21) |  |  |  |  |  |  | 1.00 | <0.01 |
| Bis+Zp+Pol | 1.50 (1.54) |  |  |  |  |  |  |  | <0.01 |
| No treatment | 5.65 (2.83) |  |  |  |  |  |  |  |  |
|  | n=237 |  |  |  |  |  |  |  |  |

**Table 4:** Clinical severity scores at 48 hours post-infection in a murine *P. aeruginosa* corneal infection model. Clinical scores were assessed 48 h post-infection in the same experimental cohort described in Table 1. Each image was independently graded twice by two blinded observers (Cohen’s weighted κ = 0.74–0.84, indicating strong inter- and intra-rater agreement). Data represent mean ± SD. Statistical analysis was performed using a General Linear Model (GLM) with Tukey’s post hoc test (*p* < 0.05). Compared to mock-treated infected controls, combination treatments (Bis + Pol, Zp + Pol, Bis + Zp + Pol) significantly improved clinical outcome, achieving scores comparable to uninfected controls.

| 48 h | Mean (SD)<br>Clinical<br>score | Condition |  |  |  |  |  |  |  |
| --- | --- | --- | --- | --- | --- | --- | --- | --- | --- |
| Condition |  | Bis | Zp | Pol | Bis+Zp | Bis+Pol | Zp+Pol | Bis+Zp+Pol | Infection |
| No infection | 0.80 (0.96) | p<0.01 | <0.01 | <0.01 | <0.01 | 0.95 | 0.46 | 0.13 | <0.01 |
| Bis | 9.05 (5.27) |  | 0.99 | 0.87 | 0.90 | 0.003 | 0.26 | 0.17 | 0.97 |
| Zp | 7.50 (5.48) |  |  | 1.00 | 1.00 | 0.06 | 0.78 | 0.73 | 0.25 |
| Pol | 6.91 (5.00) |  |  |  | 1.00 | 0.10 | 0.91 | 0.88 | 0.03 |
| Bis+Zp | 7.09 (5.50) |  |  |  |  | 0.053 | 0.84 | 0.79 | <0.03 |
| Bis+Pol | 2.67 (3.13) |  |  |  |  |  | 0.98 | 0.89 | <0.01 |
| Zp+Pol | 4.55 (3.61) |  |  |  |  |  |  | 1.00 | 0.005 |
| Bis+Zp+Pol | 4.88 (4.86) |  |  |  |  |  |  |  | <0.01 |
| No treatment | 10.53 (4.40) |  |  |  |  |  |  |  |  |
|  | n=209 |  |  |  |  |  |  |  |  |

Combinations of the compounds showed lower clinical scores on both days 1 and 2 compared to individual compounds (tables 3 and 4). Combinations were all significantly different to infection by day 2 but were not significantly different to no infection.

Non-infected corneas exhibited minimal pathology, with low clinical scores. Infected corneas receiving mock treatment (PBS with 0.1% (v/v) DMSO) displayed substantial disease, with a mean clinical score of 10.53 (Figure 9A). Treatment with individual ExoU inhibitors Bis, Zp or Pol, did not significantly reduce clinical scores compared to mock. In contrast, combinatorial treatments led to marked improvement (Figures 9A and B). Bis+Pol and Zp+Pol significantly reduced clinical scores, with Bis+Pol showing the greatest benefit. The triple treatment (Bis+Zp+Pol) also improved outcomes relative to mock treatment, though it was not significantly different from the most effective double combinations (Table 4).

Despite these clinical improvements, bacterial load analysis at 48 hours revealed no statistically significant differences in *P. aeruginosa* CFU across treatment groups (Figure 9C), indicating that the observed therapeutic benefit was not due to reduced bacterial burden.

### Assessment of immune cell viability and corneal infiltration

Flow cytometric analysis revealed comparable numbers of total non-immune (CD45⁻) cells across all treatment groups, indicating similar recovery of total corneal cells per sample following infection and treatment (Figure 9D). However, assessment of cell viability showed that combination treatments with Bis+Zp, Zp+Pol, and Bis+Zp+Pol significantly reduced the proportion of dead CD45⁻ cells compared to the mock-treated group (Figure 9E). No such reduction was observed with single-compound treatments. Analysis of immune cell infiltration revealed a significant reduction in total neutrophil numbers (CD45⁺Ly6B⁺F4/80⁻) only in the Bis+Pol and Zp+Pol treatment groups (Figure 9F). Furthermore, the proportion of dead neutrophils was significantly decreased in the Bis+Pol and Bis+Zp+Pol groups, suggesting that these treatments provided a protective effect on infiltrating neutrophils (Figure 9G).

## Discussion

Microbial keratitis, as the second most common cause of blindness remains an important clinical problem [40], with ExoU producing strains of *P. aeruginosa* being a major component of the disease [8]. Targeting the phospholipase activity of ExoU in the host cell would be a reasonable strategy to overcome the documented partial clinical efficacy of antimicrobials [8, 22]. In this study we incorporated further steps into our pipeline for the discovery of ExoU inhibitors. The first high-throughput real-time phospholipase assay [22] was followed by a high content co-infection assay of *P. aeruginosa* cytotoxicity using the Incucyte and the CD7 platforms, which were more robust and higher throughput than the manual scratch wound assay use previously [22]. These were complemented by measurement of compound inhibitory action on the *ex vivo* porcine corneal model, on pathogen lethality in *Galleria* and on a murine model of microbial keratitis. This integrated pipeline, combining in vitro and cell-based analyses of ExoU inhibition, enabled the identification of Bis, Pol, and Zp as safe and effective inhibitors that significantly reduced ExoU-mediated corneal damage in mice. Moreover, the counter-screens against human PLA₂G7 and PLA₂G4C (Supplementary Figure 5), demonstrated their ExoU selectivity, which reduces concerns over collateral impairment of host lipid signalling or membrane homeostasis, as found for some phospholipase inhibitors, such as darapladib [41].

The analysis of the likely pharmacophores of the three compounds demonstrated that whereas the metal cations Zn^2+^ was active an inhibitor *in vitro* (Figures 2B, C) in the HCT-E cell co-infection assays ZnSO_4_ was without activity, presumably because they either did not have access to the cell interior or were incorporated into the cell’s metal ion metabolism. In both cases they would be unable to exert effects on the ExoU secreted into host cells by *P. aeruginosa*’s T3SS. Thus, a chelate of Zn^2+^ ions was required, though the chelator pyrithione had no effect on ExoU catalytic activity. Given ExoU’s dependence on a catalytic serine [42], we speculate that Zn²⁺ may coordinate key active site residues, thereby inactivating the enzyme. Bismuth’s unique thiophilicity and propensity to form stable, multidentate complexes [43] suggest a mechanism distinct from zinc (Figure 3). Bi³⁺ only induced significant perturbation of ExoU thermal stability in the presence of lipid, suggesting an allosteric mechanism whereby Bi³⁺ preferentially targets the activated, membrane-bound conformation (Figure 3A). Structural studies, such as zinc/bismuth-soak co-crystallography, would enable the binding site of these metal ions to be established and to determine if the chelate transfers to metal cation to the ExoU or if the remaining coordination sites in the chelate are sufficient for binding. Structurally, Pol mimics a phospholipid: its hydrophobic acyl tail resembles a fatty acyl chain, and at the sn-2 position, normally a double O-C-P linkage in native phospholipids, it instead features a double O-N bond to diaminobutyric acid (Dab), likely preserving key electrostatic and spatial cues. It appeared that peptide length, charge complementarity and lipid anchoring were essential for its ExoU inhibitory activity. The requirement for both the cyclic peptide and fatty acyl tail suggested that Pol simultaneously occupied ExoU’s phospholipid-binding groove and an adjacent hydrophobic surface, sterically blocking substrate access while destabilising the enzyme’s active conformation. To delineate these interactions at high resolution, approaches such as site-specific photo-crosslinking or hydrogen-deuterium exchange mass spectrometry could identify critical “hotspots” within ExoU [44]. The insights gained could then guide the design of smaller mimetics that maintain both modes of engagement blocking catalysis and disfavouring the active fold without the known nephrotoxic liabilities of polymyxins [45].

Thus, although detailed structure-function data are not available, the present data demonstrate that the two metal cations and the Pol amphipathic peptide likely have three distinct modes of inhibition, zinc-mediated active-site blockade, bismuth-driven allosteric destabilisation, and dual-component peptide-lipid engagement. This provided an important mechanistic framework for rational optimisation of ExoU inhibitors as a multicomponent formulation.

In HCE-T cells Bis was not toxic at the concentrations tested but Zp and Pol had demonstrable toxicity in HCE-T, with LD_50_ of 33 µM and 43 µM, respectively (Supplementary Figure 4). For the cell infection assays the formulations tested were all well below these toxic concentrations. Moreover, to avoid the results being confounded by the antimicrobial activity of Pol, its concentration was below its MIC. There are considerable human safety data available on all three compounds. Zinc salts, particularly zinc sulphate, have long been used in ophthalmic formulations as mild astringents or antiseptics to relieve irritation and conjunctival congestion [46, 47]. Commercial products containing zinc sulphate or zinc gluconate are marketed in several countries, often in combination with vasoconstrictors or soothing excipients. Bismuth containing ointments, such as bibrocatherol, are currently used in the management of eyelid inflammation [48] and polymyxinDB is formulated in eye drops to treat microbial keratitis [49]. In mice the concentration of Zp in the formulation was raised to 20 µM (Figure 9), which is close to its 33 µM LD_50_ in HCE-T cells. Taken together, thus, although the safety of a formulation of all three compounds remains to be established, individually they could certainly be used at concentrations considerably higher than in the present cell formulations and it is likely that a formulation of all three compounds at higher concentrations would be safe in the eye.

The complementary mechanisms of action of the three compounds, demonstrated by analysis of their activity *in vitro* (Figure 3) and their effects on EGFP-ExoU(S142A) in cells (Figure 4) supported the use of the compounds in combination. Moreover, the degradation of EGFP-ExoU(S142A) induced by Bis and Pol (Figure 4) suggests these compounds may afford two layers of protection, direct inhibition of the enzyme and its elimination. It may be that Bis and Pol amplify the known routing to ExoU to vacuolar structures in cells [35, 36] potentially by altering ExoU’s conformation or membrane interactions [50] in a way that exposes lysosomal targeting signals or disrupts interactions with stabilising cofactors [51]. However, it cannot be excluded that off-target effects of Bis and Pol cause a general upregulation of autophagy in the cells.

The analysis of the protection afforded by the individual compounds and two of their pairs against a panel of 52 clinical *P. aeruginosa* keratitis isolates provided important insight into the translational robustness and limitations of these compounds as an ExoU-targeted therapy. While the laboratory strains PA103 offered a controlled proof-of-concept, clinical isolates encompass the genetic and phenotypic diversity encountered in patients [37]. The high-content microscopy screen demonstrated first that the compounds were selective, as they had little effect on the 26 exoS^+^ strains. Then, they demonstrated that whereas single-agent treatments protected cells against 48% to 52% of ExoU-positive strains (Figures 6B, C and D), the pairs (Bis+Zp, Bis+Pol) were more effective, increasing protection to 62-66% of *ExoU^+^* strains.

In the porcine corneal infection model the Bis, Zp, nor Pol monotherapies did not fully prevent PA103-induced opacity or ulceration, whereas combination treatments, particularly Bis+Pol and the triple regimen Bis+Zp+Pol, markedly improved corneal clarity and epithelial integrity, achieving near-complete protection comparable to high-dose moxifloxacin (Figure 7). The absence of significant reductions in bacterial burden demonstrated that these inhibitors acted through anti-virulence rather than bactericidal mechanisms, so preserving host tissue without driving pathogen clearance. This decoupling of therapeutic effect from antimicrobial activity is likely advantageous, as it may avoid selective pressures that contribute to antibiotic resistance [52]. Furthermore, histological preservation of the epithelium and stroma in treated corneas highlights the clinical relevance of ExoU neutralisation (Figure 7E): by maintaining barrier function, these compounds could reduce patient morbidity, accelerate healing, and prevent secondary infections.

The *Galleria mellonella* infection studies provided an *in vivo* proof-of-concept for ExoU inhibitor efficacy in a living host (Figure 8). Monotherapies with Bis, Zp, or Pol each only modestly delayed larval mortality, combination regimens, particularly Bis+Pol and the triple Bis+Zp+Pol cocktail significantly prolonged survival and health scores (Figure 8). These findings mirror our *ex vivo* corneal data and underscored the advantage of a therapeutic formulation consisting of all three compounds.

The murine scratch-and-infection model of *P. aeruginosa* keratitis provided key elements of the immune response, tear dynamics, and *in vivo* pharmacokinetics [24]. Consistent with *ex vivo* and *Galleria* results, individual treatments with Bis, Zp, or Pol failed to significantly reduce clinical scores or improve corneal pathology (Figure 9A, Tables 2 and 3). In contrast, combination regimens, particularly Bis+Pol and Zp+Pol, yielded marked reductions in clinical severity by 48 h, approaching the near-normal scores observed in uninfected controls. The triple combination (Bis+Zp+Pol) further enhanced corneal protection, albeit without statistical advantage over the best dual therapies, suggesting a plateau of therapeutic benefit. Again, bacterial CFU counts remained unchanged across all inhibitor conditions, confirming an anti-virulence rather than bactericidal mechanism (Figure 9C). Flow cytometry of digested corneas revealed that dual and triple inhibitor treatments preserved viability of epithelial cells included in the CD45⁻ population and reduced neutrophil infiltration and death, an indication that inhibition of ExoU-mediated cytotoxicity helped maintain both barrier integrity and innate immune cell function (Figure 9D-G). The sparing of neutrophils, which play a key role in bacterial clearance and tissue repair, may further contribute to improved clinical outcomes independent of direct bacterial killing.

## Conclusions

Across biochemical, cellular, *ex vivo*, and *in vivo* models, we demonstrate that repurposed FDA-approved agents zinc pyrithione, bismuth subcitrate, and polymyxinDB, effectively inhibit the *P. aeruginosa* virulence factor ExoU by distinct but complementary mechanisms. Monotherapies provided partial protection, while dual and triple combinations delivered considerable improvements in enzyme inhibition, cell viability, tissue integrity, and host survival. Importantly, these anti-virulence effects occur without reducing bacterial burden or inhibiting T3SS gene expression, highlighting a strategy to neutralise pathogenicity without driving antibiotic resistance. Our work establishes a robust framework for combination anti-ExoU therapies and paves the way for clinical translation to improve outcomes in bacterial keratitis and highlights the potential of anti-virulence strategies as complements to existing antibiotics and reducing the risk of resistance development [53]. By recapitulating a principle well-established in antimicrobial chemotherapy, where drug combinations delay resistance emergence [54], anti-virulence combinations may similarly forestall adaptive changes in *P. aeruginosa* that could reduce monotherapy effectiveness.

## Acknowledgments

We acknowledge the Centre for Cell Imaging (CCI) of the University of Liverpool Shared Research Facilities, and the Cleveland Clinic Research Flow Cytometry Core, for provision of imaging equipment and flow cytometer respectively, as well as technical assistance. We would specifically like to thank Thomas Waring for technical assistance. The Zeiss Cell Discoverer 7 microscope was purchased through grant MR/X013502/1 and the LSM 780 was funded by MRC grant number MR/K015931/1.

## Supplementary Figures

**Supplementary Figure 1.**
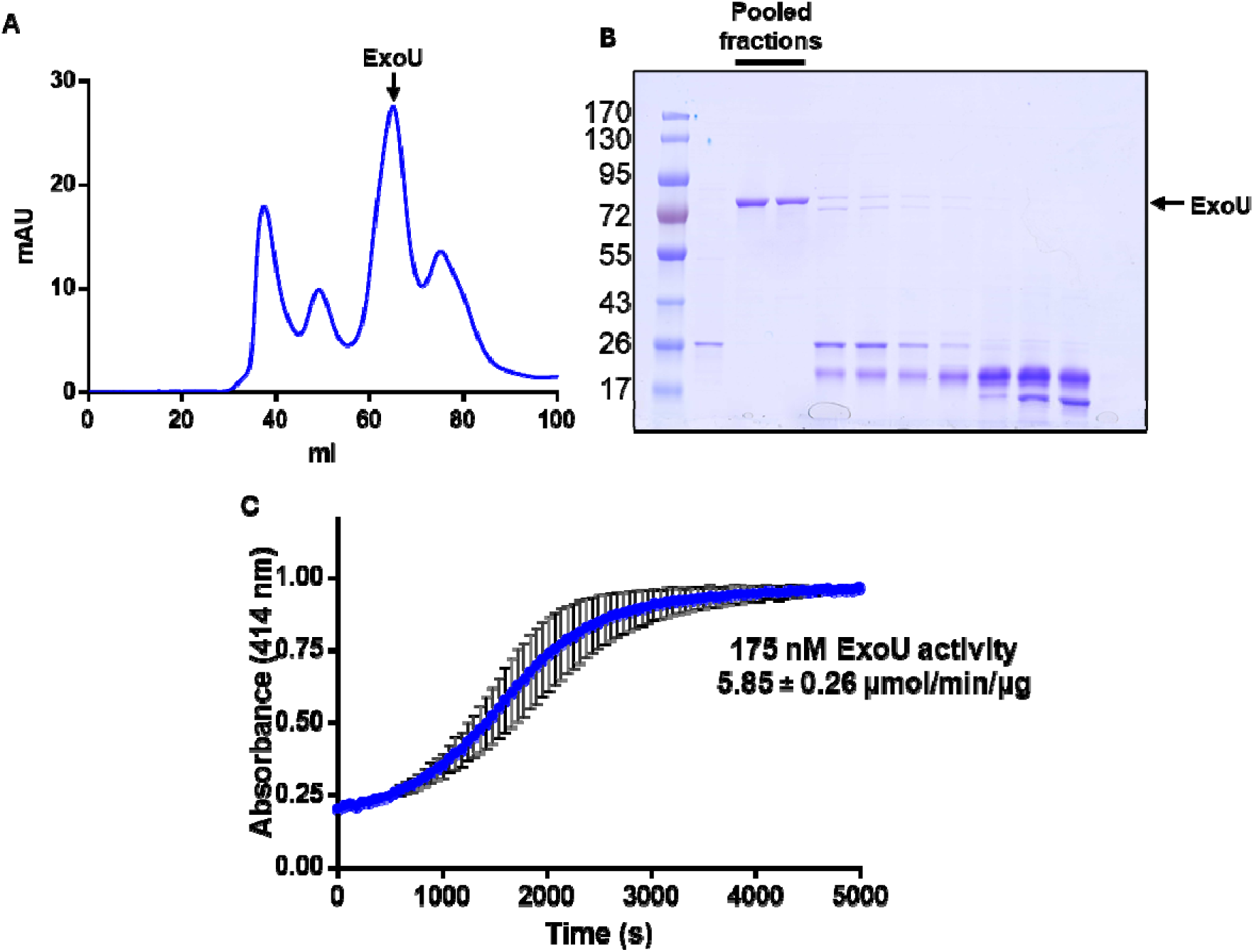
Purification of and analysis of ExoU phospholipase activity: His-tagged ExoU was purified from C43(DE3) E. coli by Immobilised metal affinity chromatography (IMAC) followed by (A) size exclusion chromatography prior to (B) SDS-PAGE analysis. (C) ExoU hydrolysis of arachidonoyl Thio-PC substrate was analysed in the presence of activating co-factors PIP_2_ (0.5 µM) and mono-ubiquitin (10 µM).

**Supplementary Figure 2.**
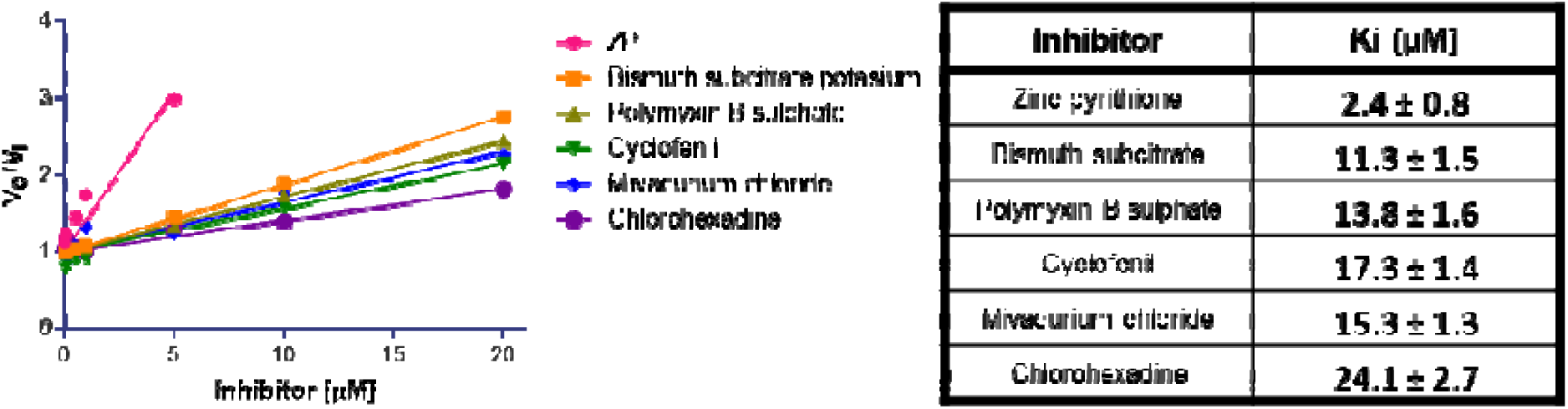
Dose-response analysis to determine ExoU inhibitor Ki values. Phospholipase assays were performed with varying concentrations of identified inhibitors. Inhibition at each inhibitor concentration was plotted against log[inhibitor] and fitted to a four-parameter logistic model to obtain IC₅₀ values. Ki values were calculated using the Cheng–Prusoff equation, incorporating the independently determined Km for ExoU and the substrate concentration used in the assay. Each data point represents the mean ± SD of three independent experiments, each performed in triplicate. Curve fitting and Ki calculations were performed in GraphPad Prism 9.0.

**Supplementary Figure 3.**
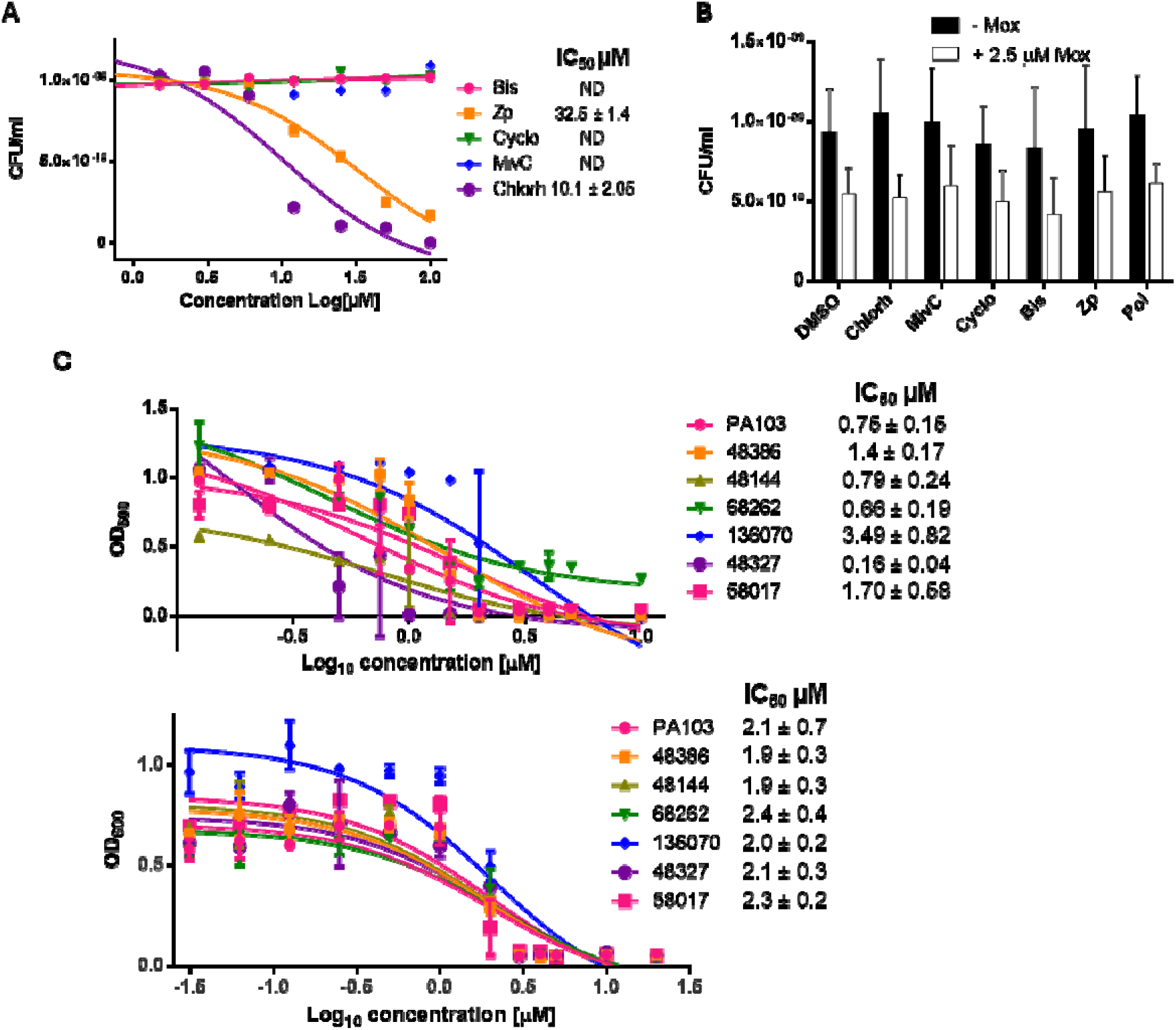
ExoU inhibitors are not antimicrobial at concentrations used in cellular assays: (A) Dose-response growth curves of *P*. *aeruginosa* PA103 in LB broth treated with candidate inhibitors, measured by endpoint CFU enumeration after overnight incubation.(B) Endpoint CFU enumeration of *P. aeruginosa* PA103 cultures collected following IncuCyte S3 automated time-lapse microscopy analysis to monitor bacterial growth dynamics in with and without 2.5 µM moxifloxacin. (C) Dose-response growth inhibition of PA103 and a panel of clinical keratitis isolates treated with Pol, determined by OD₆₀₀ measurements after overnight incubation in LB broth (top) and DMEM-F12 cell culture medium (bottom).

**Supplementary Figure 4.**
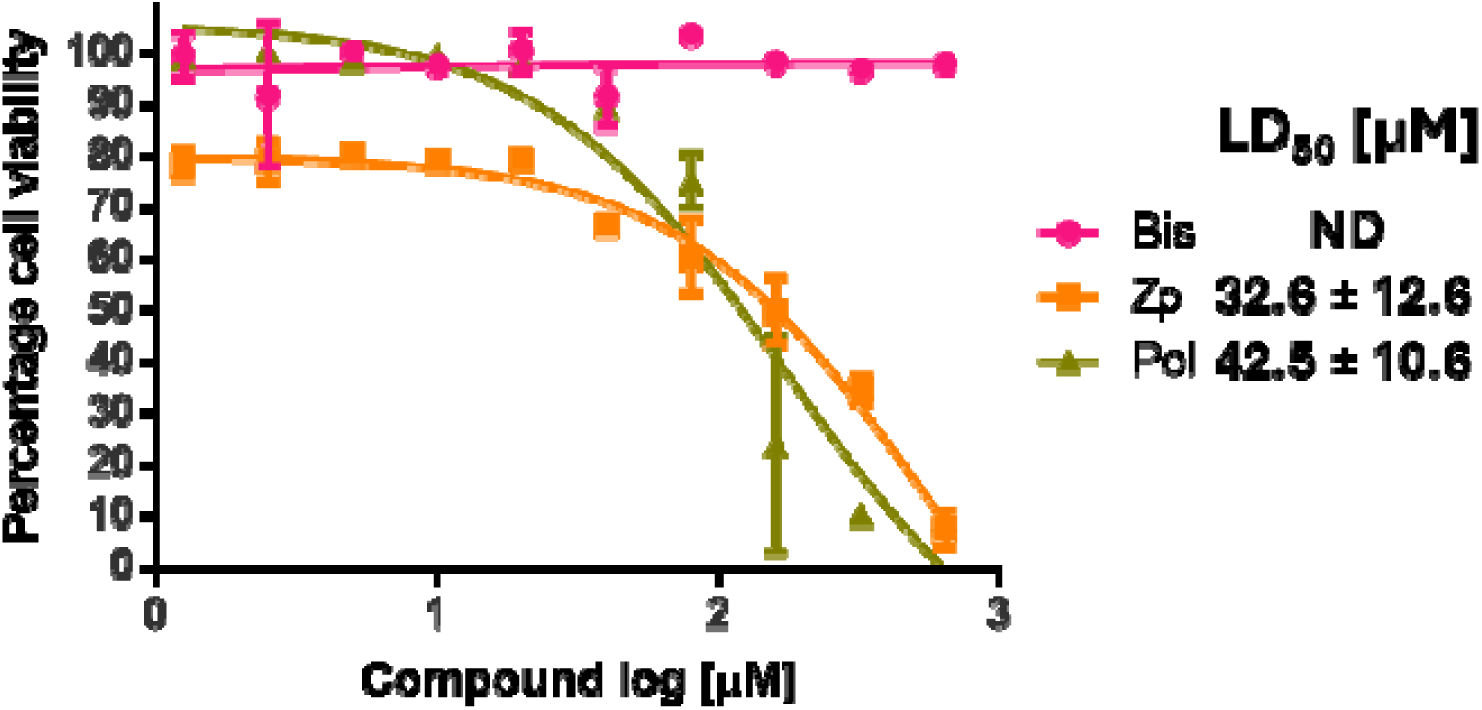
Toxicity analysis of ExoU inhibitors in human primary corneal cells: Lactate dehydrogenase (LDH) release was measured after 48 h of compound exposure to assess host cell compatibility. Cells were seeded at 8,000 cells/well, treated with compounds at the indicated concentrations, and LDH release was quantified using the Cytotoxicity Detection Kit (Roche). Values were background-subtracted, normalized to maximal LDH release controls, and expressed as mean ± SD (n = 3 independent experiments, each in duplicate). Nonlinear regression analysis (variable slope) was used to calculate LD₅₀ values.

**Supplementary Figure 5.**
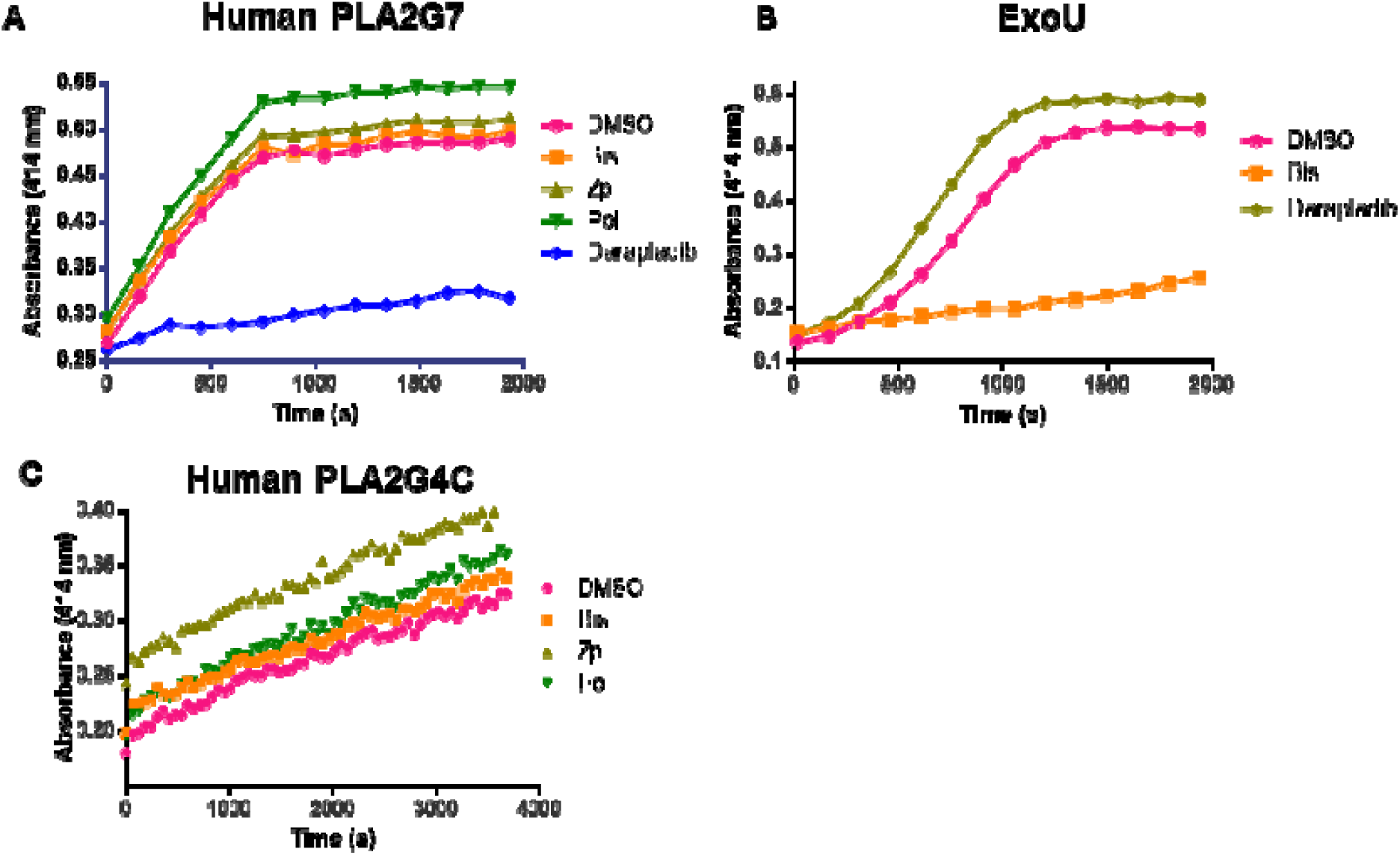
Bis, Zp and Pol do not inhibit human PLA2 enzymes: (A) Activity of human lipoprotein-associated phospholipase A₂ (PLA2G7) was measured using 2-thio-PAF as a substrate in the presence of 10 µM indicated inhibitor or 2% (v/v) DMSO controls. (B) ExoU activity was assessed using arachidonoyl thio-phosphatidylcholine (thio-PC) in the presence of 10 µM of the indicated compound. (C) Activity of human cytosolic phospholipase A₂ (PLA2G4C) was measured using arachidonoyl thio-phosphatidylcholine (thio-BGPC) in the presence of Bis, Pol, or Zp (10 µM).

**Supplementary Figure 6.**
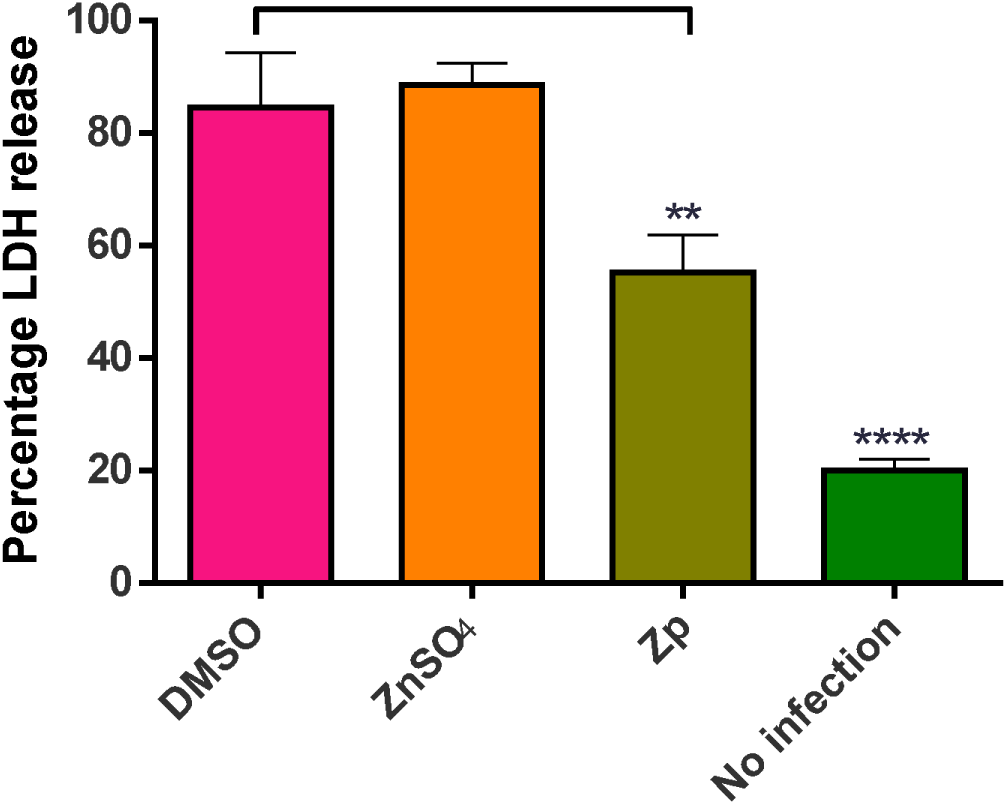
ZnSO4 does not protect HCE-T cells from ExoU cell lysis during *in vitro* PA103 infection: HCE-T cells were infected with PA103 (MOI 10) for 4 hours in the presence of either DMSO (0.1% v/v), 2.5 µM ZnSO_4_ or 2.5 µM Zp, followed by LDH assay analysis to detect cell lysis. Percentage lysis was normalized to the maximal LDH release from the kit positive control (CyQUANT™; Thermo Fisher).

**Supplementary Figure 7.**
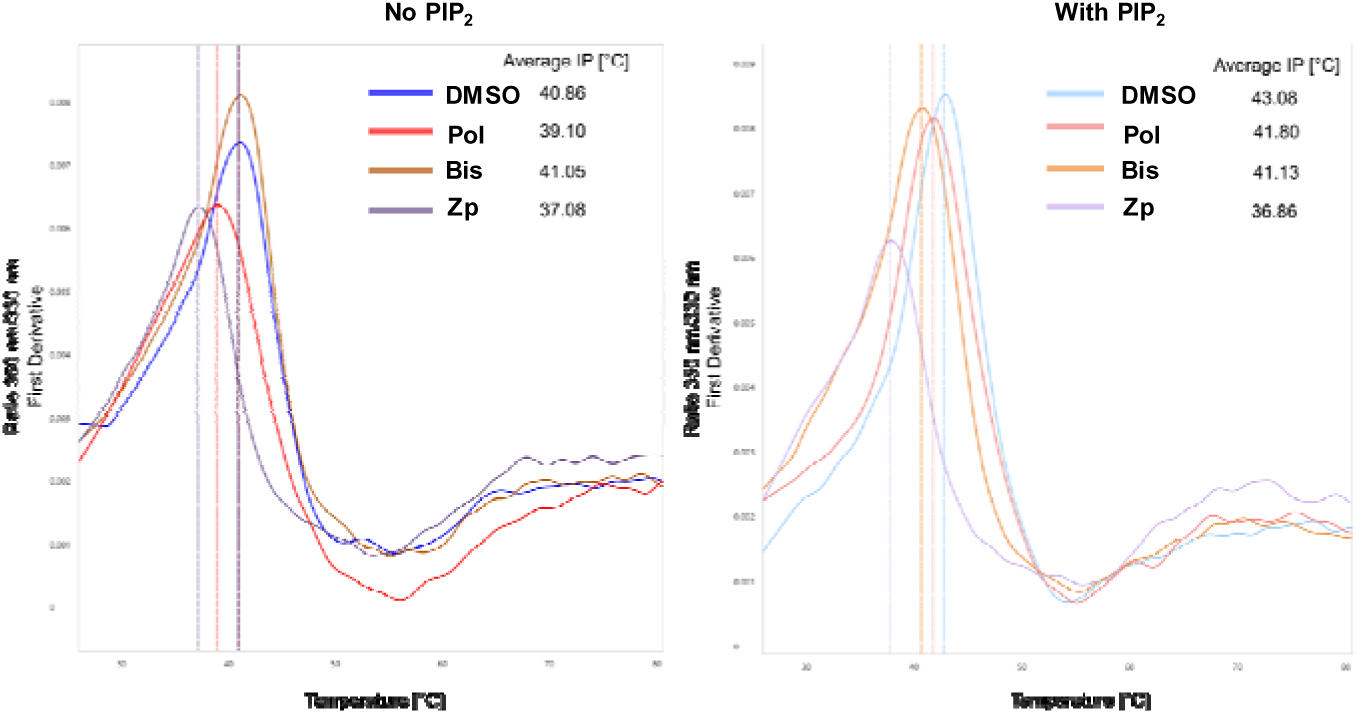
Nano differential scanning fluorimetry (nanoDSF) analysis of ExoU thermal stability in the presence of compounds. Raw melt curves of recombinant ExoU are shown in the absence (left) and presence (right) of PIP₂ (5 µM). Compounds were tested at 10 µM. Intrinsic tryptophan fluorescence emission was continuously monitored at 330 nm and 350 nm as the temperature increased, and the fluorescence ratio (F₃₅₀/F₃₃₀) was plotted against temperature. Melting temperatures (Tl7) were calculated from the average inflection point (IP) of the first derivative of the fluorescence ratio using the manufacturer’s analysis software.

**Supplementary Figure 8.**
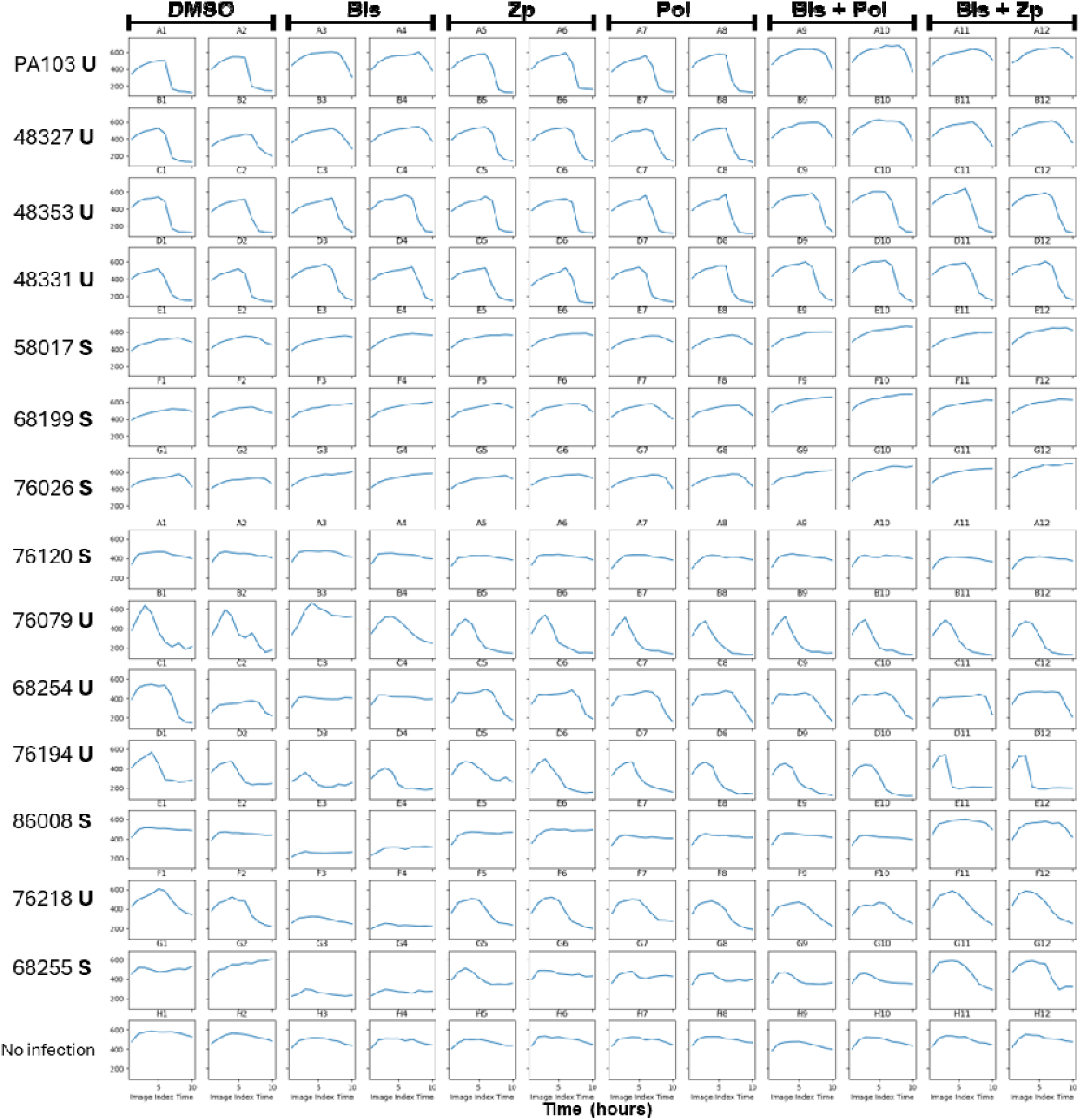
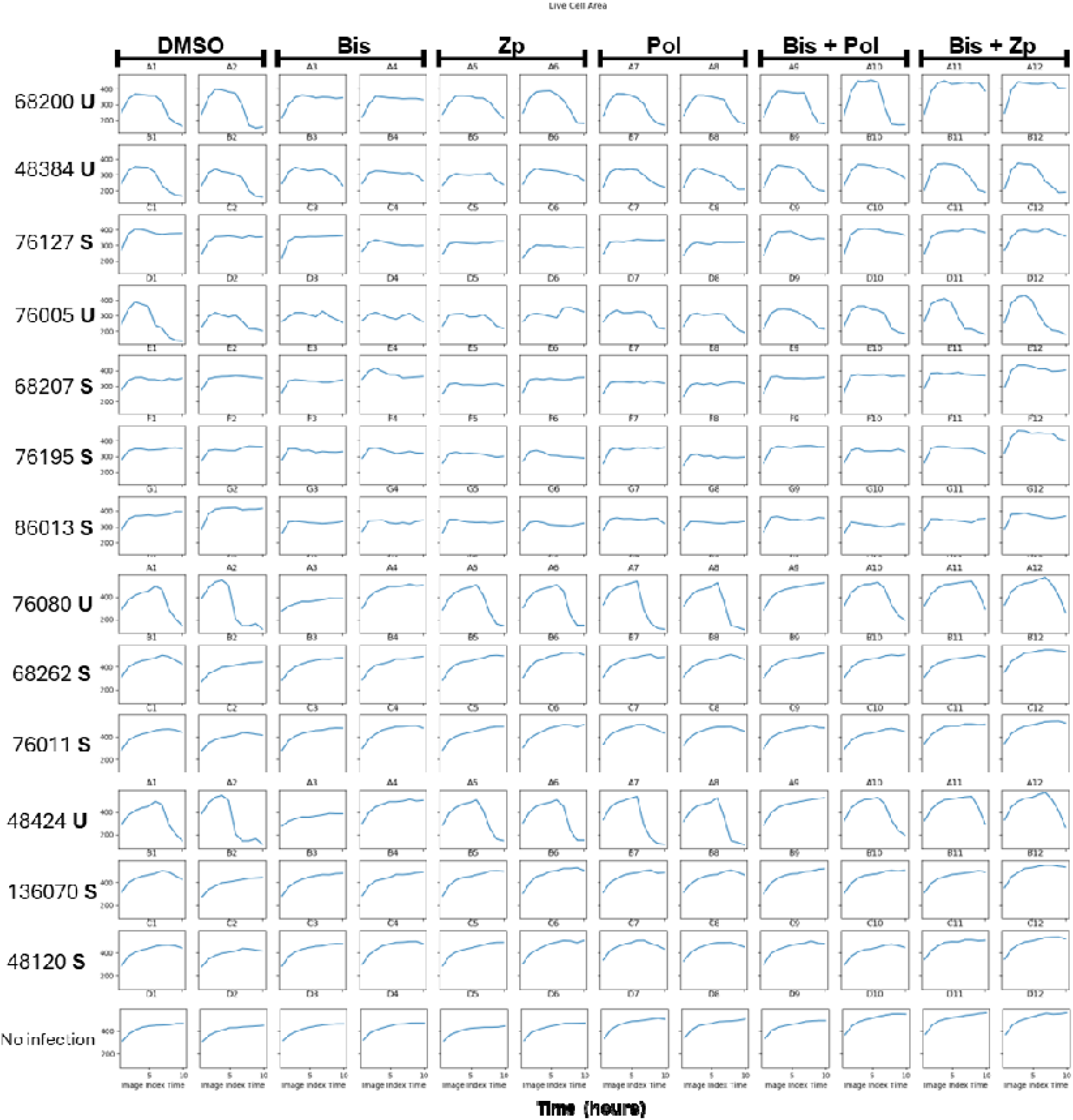
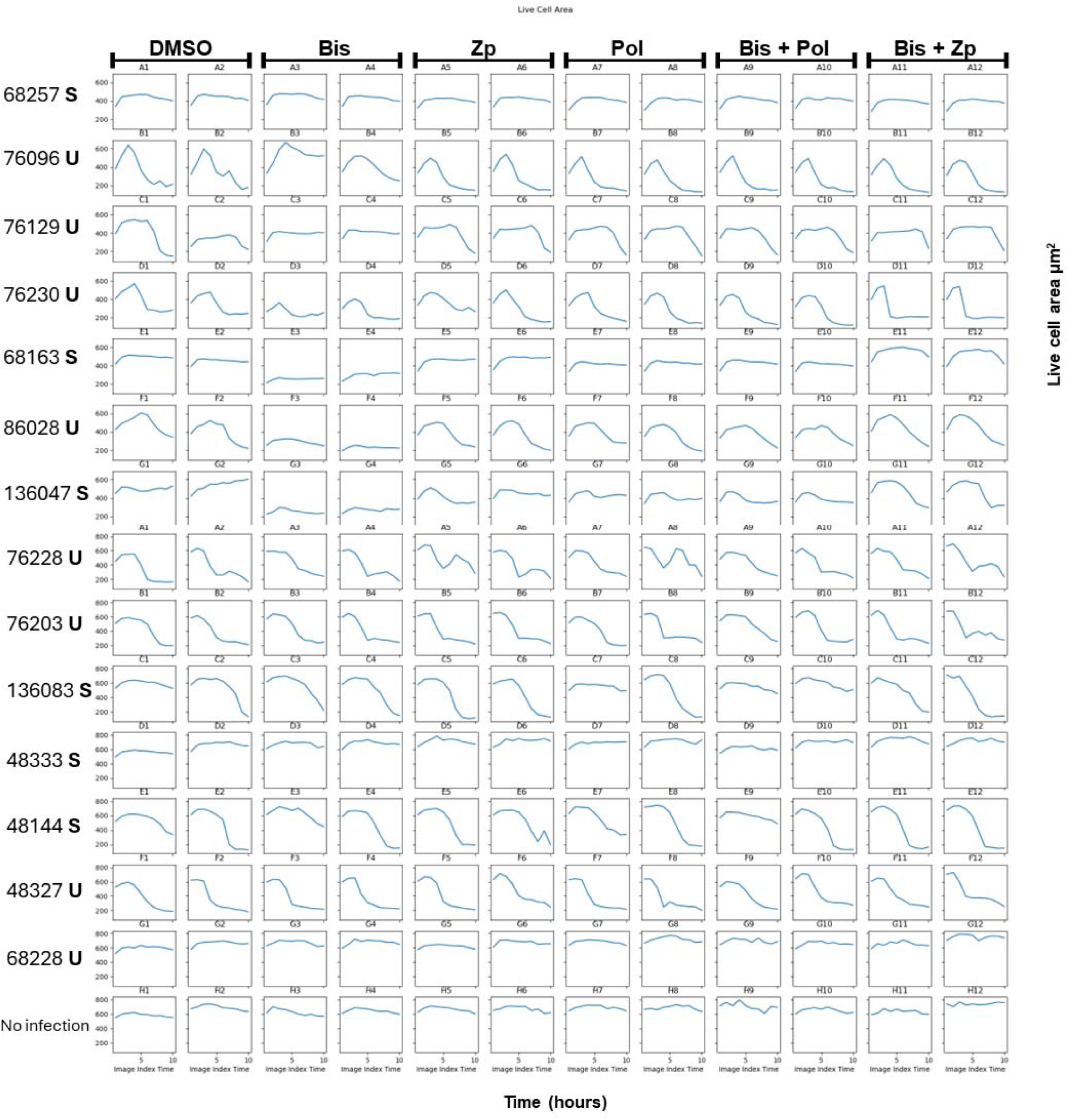
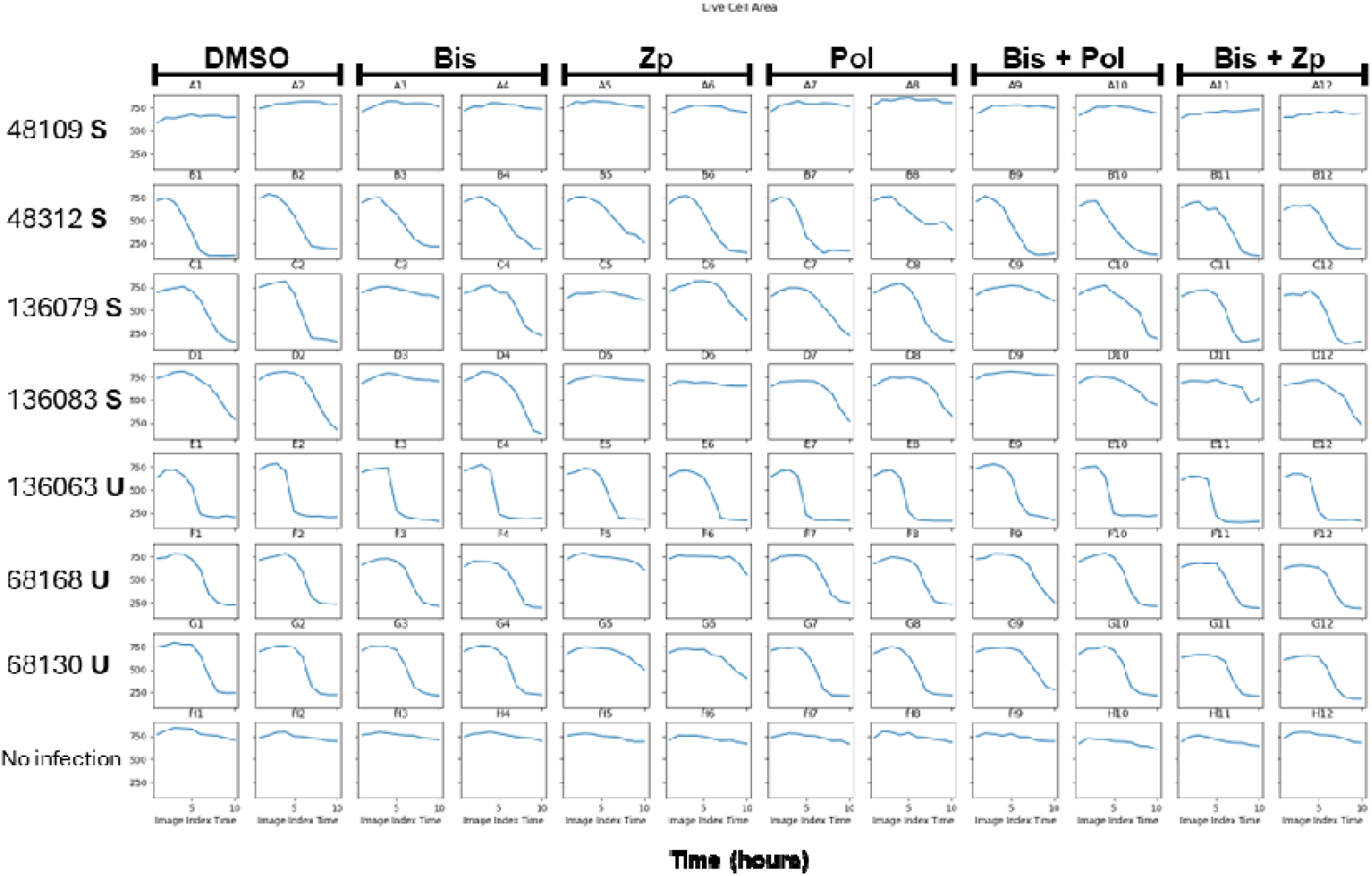
Time-course of live-cell area for all clinical keratitis isolates. Live-cell area (µm²) was measured hourly over 10 h for each *P. aeruginosa* keratitis isolate using the Cell Discover_J7 high-content imaging pipeline described in Figure_J6. Each row corresponds to a single strain, labelled on the left with “U” (ExoU-positive) or “S” (ExoS-positive). Columns represent treatment conditions, 10 µM bismuth subcitrate, 2.5_JµM zinc pyrithione, 1 µM polymyxin_JB, Bis_J+_JPol, and Bis_J+_JZp, as indicated at the top.

**Supplementary Figure 9.**
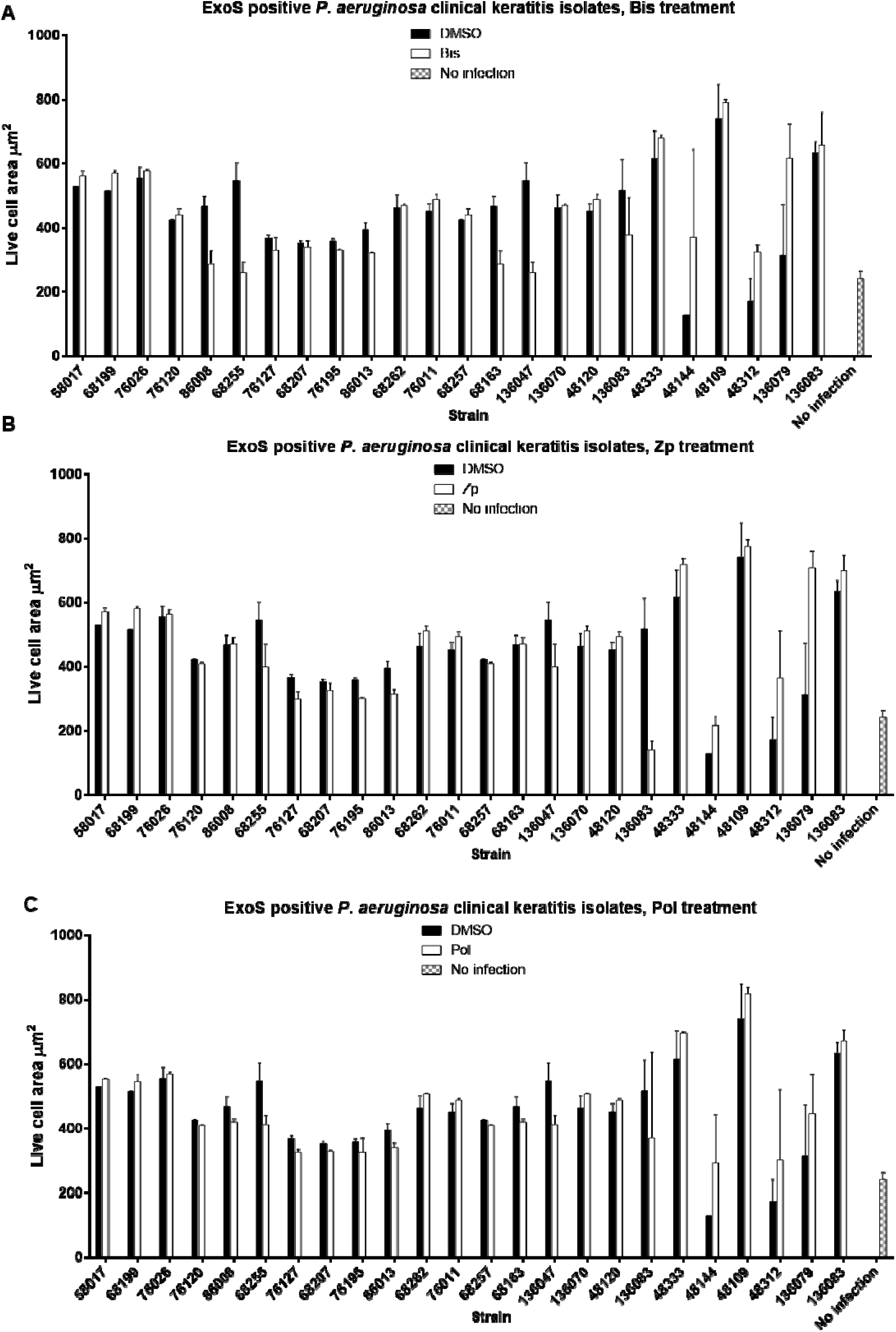
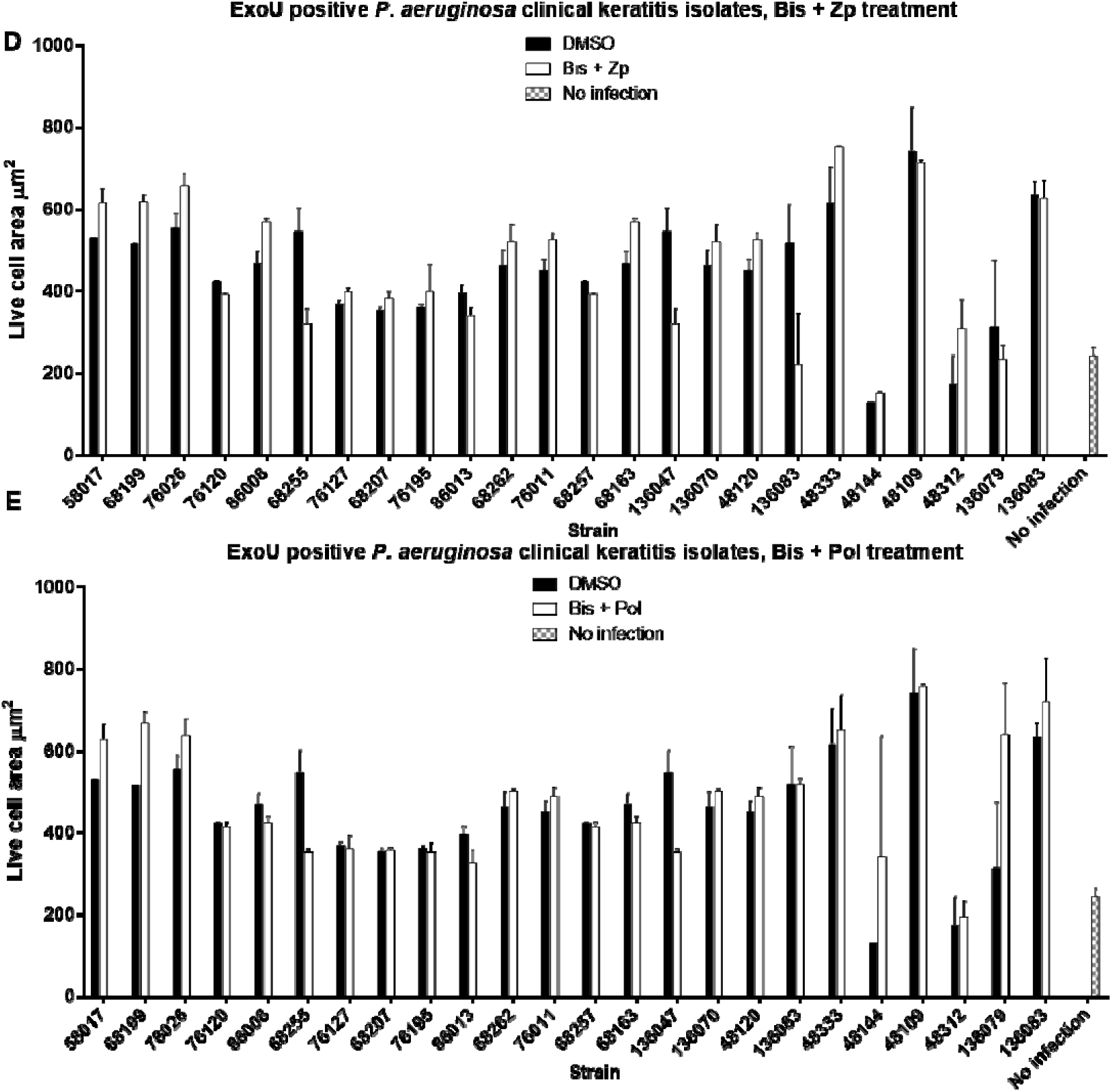
Cell convexity in HCE-T cells infected with ExoS-positive *P. aeruginosa* isolates. Cell convexity was quantified at 8 h post-infection for 24 ExoS-expressing clinical keratitis strains using the same high-content imaging and analysis pipeline described in Figure_J6 (see legend and main text). Convexity metrics were calculated in ZEN Blue and plotted via the automated Python workflow. Bars represent mean convexity for each isolate under the following treatments: Bis 10 µM, Zp, 2.5 µM and Pol 1 µM.

**Supplementary Figure 10.**
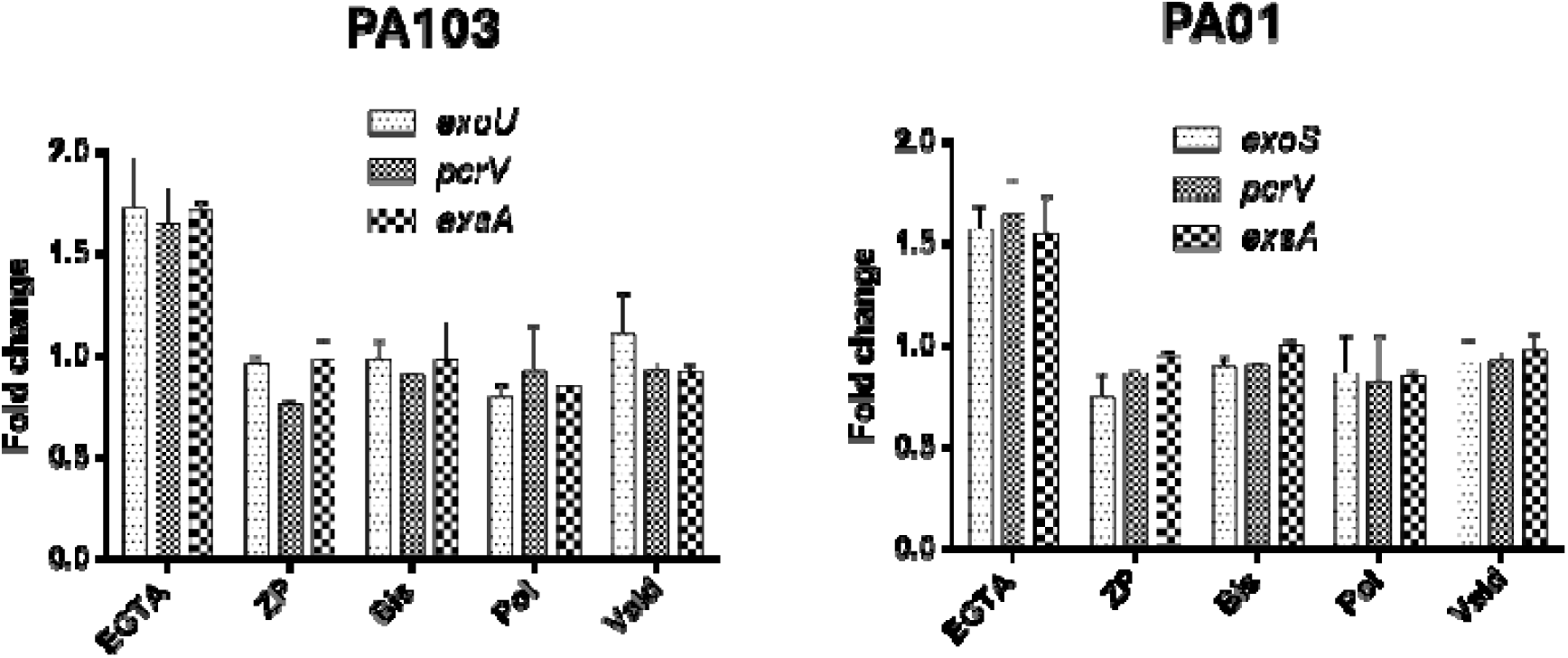
Quantitative RT-PCR analysis of Type III Secretion System (T3SS) gene expression. Gene expression of key T3SS components was assessed following bacterial growth in LB medium supplemented with bismuth subcitrate (Bis, 10 µM), zinc pyrithione (Zp, 2.5_JµM), or polymyxin B (Pol, 1 µM). For *P. aeruginosa* PA103, *exsA* (T3SS master regulator), *pcrV* (translocon component), and *exoU* (effector) were analysed; for PAO1, *exsA*, *pcrV*, and *exoS* were examined. Expression levels were normalised to RNA polymerase β-subunit (*rpoB*). EGTA (2.5_JmM) treatment served as a positive control to induce T3SS expression.

---

**Supplementary Table 1:** Results of one-way ANOVA with Tukey’s post hoc multiple comparisons performed in SPSS. The analysis was conducted on mean live cell area values averaged across all timepoints in each time series of infections with ExoU-producing *P. aeruginosa* isolates (Figure 6A). The table reports mean differences between treatment groups (D=DMSO, B=Bis, Z=Zp, P=Pol, BZ=Bis+Zp, BP=Bis+Pol), standard error, significance values, and the lower and upper bounds of the 95% confidence intervals.

| (I) Compounds | (J) Compounds | Mean Difference (I-J) | Std. Error | Sig. | 95% Confidence Interval |  |
| --- | --- | --- | --- | --- | --- | --- |
|  |  |  |  |  | Lower Bound | Upper Bound |
| B | BP | -10.9574* | 3.17739 | .008 | -20.0247 | -1.8900 |
|  | BZ | -17.7333* | 3.17739 | <.001 | -26.8006 | -8.6659 |
|  | D | 20.6243* | 3.17739 | <.001 | 11.5569 | 29.6917 |
|  | P | 25.9292* | 3.17739 | <.001 | 16.8618 | 34.9966 |
|  | Z | .6858 | 3.17739 | 1.000 | -8.3816 | 9.7532 |
| BP | B | 10.9574* | 3.17739 | .008 | 1.8900 | 20.0247 |
|  | BZ | -6.7759 | 3.17739 | .271 | -15.8433 | 2.2915 |
|  | D | 31.5817* | 3.17739 | <.001 | 22.5143 | 40.6491 |
|  | P | 36.8866* | 3.17739 | <.001 | 27.8192 | 45.9539 |
|  | Z | 11.6431* | 3.17739 | .004 | 2.5757 | 20.7105 |
| BZ | B | 17.7333* | 3.17739 | <.001 | 8.6659 | 26.8006 |
|  | BP | 6.7759 | 3.17739 | .271 | -2.2915 | 15.8433 |
|  | D | 38.3576* | 3.17739 | <.001 | 29.2902 | 47.4250 |
|  | P | 43.6625* | 3.17739 | <.001 | 34.5951 | 52.7298 |
|  | Z | 18.4190* | 3.17739 | <.001 | 9.3516 | 27.4864 |
| D | B | -20.6243* | 3.17739 | <.001 | -29.6917 | -11.5569 |
|  | BP | -31.5817* | 3.17739 | <.001 | -40.6491 | -22.5143 |
|  | BZ | -38.3576* | 3.17739 | <.001 | -47.4250 | -29.2902 |
|  | P | 5.3049 | 3.17739 | .552 | -3.7625 | 14.3723 |
|  | Z | -19.9385* | 3.17739 | <.001 | -29.0059 | -10.8711 |
| P | B | -25.9292* | 3.17739 | <.001 | -34.9966 | -16.8618 |
|  | BP | -36.8866* | 3.17739 | <.001 | -45.9539 | -27.8192 |
|  | BZ | -43.6625* | 3.17739 | <.001 | -52.7298 | -34.5951 |
|  | D | -5.3049 | 3.17739 | .552 | -14.3723 | 3.7625 |
|  | Z | -25.2434* | 3.17739 | <.001 | -34.3108 | -16.1760 |
| Z | B | -.6858 | 3.17739 | 1.000 | -9.7532 | 8.3816 |
|  | BP | -11.6431* | 3.17739 | .004 | -20.7105 | -2.5757 |
|  | BZ | -18.4190* | 3.17739 | <.001 | -27.4864 | -9.3516 |
|  | D | 19.9385* | 3.17739 | <.001 | 10.8711 | 29.0059 |
|  | P | 25.2434* | 3.17739 | <.001 | 16.1760 | 34.3108 |

Based on observed means.
The error term is Mean Square(Error) = 2322.037.
\*. The mean difference is significant at the 0.05 level.

**Supplementary Table 2.** Results of one-way ANOVA with Tukey’s post hoc multiple comparisons for non-infection controls. The analysis was conducted on mean live cell convexitivy values averaged across all timepoints for HCE-T cell cultures treated with individual compounds (Bis, Pol, Zp) or combinations (Bis+Zp, Bis+Pol) and infected with 24 *exoS^+^* isoaltes. The table reports mean differences between treatment groups, standard error, significance values, and the lower and upper bounds of the 95% confidence intervals. No significant differences were observed between groups, confirming that compounds did not effect ExoS mediated HCE-T cell convexity.

| (I) Conditions | (J) Conditions | Mean Difference (I-J) | Std. Error | Sig. | 95% Confidence Interval |  |
| --- | --- | --- | --- | --- | --- | --- |
|  |  |  |  |  | Lower Bound | Upper Bound |
| B | BP | -30.6449* | 9.67617 | .019 | -58.2373 | -3.0524 |
|  | BZ | -33.3619* | 9.68102 | .008 | -60.9682 | -5.7556 |
|  | D | -18.1580 | 9.67617 | .417 | -45.7505 | 9.4344 |
|  | P | -13.0960 | 9.67617 | .755 | -40.6885 | 14.4965 |
|  | Z | -26.1085 | 9.67617 | .076 | -53.7009 | 1.4840 |
| BP | B | 30.6449* | 9.67617 | .019 | 3.0524 | 58.2373 |
|  | BZ | -2.7171 | 9.68102 | 1.000 | -30.3234 | 24.8892 |
|  | D | 12.4868 | 9.67617 | .790 | -15.1057 | 40.0793 |
|  | P | 17.5488 | 9.67617 | .457 | -10.0436 | 45.1413 |
|  | Z | 4.5364 | 9.67617 | .997 | -23.0561 | 32.1289 |
| BZ | B | 33.3619* | 9.68102 | .008 | 5.7556 | 60.9682 |
|  | BP | 2.7171 | 9.68102 | 1.000 | -24.8892 | 30.3234 |
|  | D | 15.2039 | 9.68102 | .618 | -12.4024 | 42.8102 |
|  | P | 20.2659 | 9.68102 | .291 | -7.3404 | 47.8722 |
|  | Z | 7.2535 | 9.68102 | .976 | -20.3528 | 34.8598 |
| D | B | 18.1580 | 9.67617 | .417 | -9.4344 | 45.7505 |
|  | BP | -12.4868 | 9.67617 | .790 | -40.0793 | 15.1057 |
|  | BZ | -15.2039 | 9.68102 | .618 | -42.8102 | 12.4024 |
|  | P | 5.0620 | 9.67617 | .995 | -22.5304 | 32.6545 |
|  | Z | -7.9504 | 9.67617 | .964 | -35.5429 | 19.6421 |
| P | B | 13.0960 | 9.67617 | .755 | -14.4965 | 40.6885 |
|  | BP | -17.5488 | 9.67617 | .457 | -45.1413 | 10.0436 |
|  | BZ | -20.2659 | 9.68102 | .291 | -47.8722 | 7.3404 |
|  | D | -5.0620 | 9.67617 | .995 | -32.6545 | 22.5304 |
|  | Z | -13.0124 | 9.67617 | .760 | -40.6049 | 14.5800 |
| Z | B | 26.1085 | 9.67617 | .076 | -1.4840 | 53.7009 |
|  | BP | -4.5364 | 9.67617 | .997 | -32.1289 | 23.0561 |
|  | BZ | -7.2535 | 9.68102 | .976 | -34.8598 | 20.3528 |
|  | D | 7.9504 | 9.67617 | .964 | -19.6421 | 35.5429 |
|  | P | 13.0124 | 9.67617 | .760 | -14.5800 | 40.6049 |

Based on observed means.
The error term is Mean Square(Error) = 23407.065.
\*. The mean difference is significant at the 0.05 level.

